# The imprinted *Zdbf2* gene finely tunes feeding and growth in neonates

**DOI:** 10.1101/2021.01.27.428419

**Authors:** Juliane Glaser, Julian Iranzo, Maud Borensztein, Mattia Marinucci, Angelica Gualtieri, Colin Jouhanneau, Aurelie Teissandier, Carles Gaston-Massuet, Deborah Bourc’his

## Abstract

Genomic imprinting refers to the mono-allelic and parent-specific expression of a subset of genes. While long recognized for their role in embryonic development, imprinted genes have recently emerged as important modulators of postnatal physiology, notably through hypothalamus-driven functions. Here, using mouse models of loss, gain and parental inversion of expression, we report that the paternally expressed *Zdbf2* gene controls neonatal growth in mice, in a dose-sensitive but parent-of-origin-independent manner. We further found that *Zdbf2*-KO neonates failed to fully activate hypothalamic circuits that stimulate appetite, and suffered milk deprivation and diminished circulating Insulin Growth Factor 1 (IGF-1). Consequently, only half of *Zdbf2*-KO pups survived the first days after birth and those surviving were smaller. This study demonstrates that precise imprinted gene dosage is essential for vital physiological functions at the transition from intra- to extra-uterine life, here the adaptation to oral feeding and optimized body weight gain.

## INTRODUCTION

Genomic imprinting is the process by which a subset of genes is expressed from only one copy in a manner determined by the parental origin. In mammals, genomic imprinting arises from sex-specific patterning of DNA methylation during gametogenesis, which generates thousands of germline differentially methylated regions (gDMRs) between the oocyte and the spermatozoa. After fertilization, the vast majority of gDMRs are lost during the epigenetic reprogramming that the embryonic genome undergoes (Seah & Messerschmidt, 2017). However, some gDMRs are protected through sequence- and DNA methylation-specific recruitment of the KRAB-associated protein 1 (KAP1) complex (Li *et al*, 2008; Quenneville *et al*, 2011; Takahashi *et al*, 2019) and become fixed as imprinting control regions (ICRs). Roughly 20 ICRs maintain parent-specific DNA methylation throughout life and across all tissues in mouse and human genomes, and control the mono-allelic and parent-of-origin expression of approximately 150 imprinted genes (Schulz *et al*, 2008; Tucci *et al*, 2019).

DNA methylation-based genome-wide screens have led to the conclusion that all life-long ICRs have probably been discovered (Proudhon *et al*, 2012; Xie *et al*, 2012). However, a greater number of regions are subject to less robust forms of imprinted DNA methylation, restricted to early development or persisting in specific cell lineages only. Moreover, imprinted genes are often expressed in a tissue- or stage-specific manner (Proudhon *et al*, 2012; Monteagudo-Sánchez *et al*, 2019), adding to the spatio-temporal complexity of genomic imprinting regulation. Finally, while the vast majority of imprinted genes are conserved between mice and humans, a subset of them have acquired imprinting more recently in a species-specific manner (Bogutz *et al*, 2019). Why is reducing gene dosage important for imprinted genes, why does it occur in a parent-of-origin manner and why is it essential for specific organs in specific species are fundamental questions in mammalian development and physiology.

Imprinted genes have long-recognized roles in development and viability *in utero*, by balancing growth and resource exchanges between the placenta and the fetus. Moreover, it is increasingly clear that imprinted genes also strongly influence postnatal physiology (Peters, 2014). Neonatal growth, feeding behavior, metabolic rate and body temperature are affected by improper dosage of imprinted genes in mouse models and human imprinting disorders (Charalambous *et al*, 2014; Ferrón *et al*, 2011; Leighton *et al*, 1995; Li *et al*, 1999; Plagge *et al*, 2004; Nicholls *et al*, 1989; Buiting, 2010). Imprinting-related postnatal effects are recurrently linked to dysfunction of the hypothalamus (Ivanova & Kelsey, 2011), a key organ for orchestrating whole body homeostasis through a complex network of nuclei that produce and deliver neuropeptides to distinct targets, including the pituitary gland that in turn secretes endocrine hormones such as the growth hormone (GH). Accordingly, the hypothalamus appears as a privileged site for imprinted gene expression (Gregg *et al*, 2010). A typical illustration of such association is provided by a cluster of hypothalamic genes whose dosage is altered in Prader-Willi syndrome (PWS). PWS children present neurological and behavioral impairments in particular related to feeding, in the context of hypothalamic neuron anomalies (Swaab, 1997; Cassidy & Driscoll, 2009). In mouse models, single inactivation of the PWS-associated *Magel2* gene results in neonatal growth retardation, reduced food intake and altered metabolism (Bischof *et al*, 2007; Kozlov *et al*, 2007; Schaller *et al*, 2010). Fine-tuning hypothalamic inputs is particularly important for adapting to environmental changes, the most dramatic one for mammals being the transition from intra- to extra-uterine life at birth. Early mis-adaptation to postnatal life can have far-reaching consequences on adult health, by increasing the risk of metabolic diseases. It therefore is of the outmost importance to thoroughly document the action of imprinted genes, particularly in hypothalamic functions.

*Zdbf2* (*DBF-type zinc finger- containing protein 2*) is a paternally expressed gene with preferential expression in the brain (Kobayashi *et al*, 2009; Greenberg *et al*, 2017). It is one of the last-discovered genes with life-long and tissue-wide imprinted methylation, and conserved imprinting in mice and humans, however its function is not yet resolved (Kobayashi *et al*, 2009; Duffié *et al*, 2014). We previously characterized the complex parental regulation of the *Zdbf2* locus: it is controlled by a maternally methylated gDMR during the first week of embryogenesis, but for the rest of life, it harbors a somatic DMR (sDMR) that is paternally methylated (Proudhon *et al*, 2012; Duffié *et al*, 2014) (Figure1A). The maternal gDMR coincides with a promoter that drives paternal-specific expression of a *Long isoform of Zdbf2* (*Liz*) transcript. In the pluripotent embryo, *Liz* transcription triggers *in cis* DNA methylation at the sDMR, allowing de-repression of the canonical promoter of *Zdbf2,* located ~10kb downstream (Greenberg *et al*, 2017) (Figure1A). In fact, although specifically expressed in the embryo where it undergoes stringent multi-layered transcriptional control (Greenberg *et al*, 2019), *Liz* is dispensable for embryogenesis itself. Its sole function seems to epigenetically program expression of *Zdbf2* later in life: genetic loss-of-function of *Liz* (*Liz*-LOF) prevents methylation of the sDMR, giving rise to mice that cannot activate *Zdbf2*–despite an intact genetic sequence-in the hypothalamus and the pituitary gland and display reduced postnatal growth (Greenberg *et al*, 2017). The cause of this growth phenotype and whether it is directly linked to the function and dosage regulation of *Zdbf2* in the brain cells is unknown.

By generating loss-of-function (LOF) and gain-of-function (GOF) mouse mutants, we show here that ZDBF2 is necessary for optimal growth and survival during the nursing period, by stimulating hypothalamic food circuits immediately at birth. Moreover, our data support that the dose but not the parental origin of *Zdbf2* expression is important for its imprinted mode of action. Altogether, our study illustrates the critical function and proper dose regulation of *Zdbf2* for adaptation to postnatal life.

## RESULTS

### *Zdbf2* is expressed in the neuro-endocrine cells of the hypothalamo-pituitary axis

Besides a C2H2 zinc finger motif, the ZDBF2 protein does not contain any obvious functional domain that could inform its molecular function. To gain insights into the role of *Zdbf2*, we first examined the cellular specificity and temporal dynamics of its expression. Comprehensive datasets of adult tissues expression in mice and human suggested *Zdbf2* expression is prevalent in brain tissues and pituitary gland (biogps in mouse tissues: http://biogps.org/#goto=genereport&id=73884 and GTex in human tissues: https://gtexportal.org/home/gene/ZDBF2). Our data endorse this brain and pituitary-specific expression of *Zdbf2* in adult mice and show the highest level of expression in the hypothalamus (Figure S1A) (Greenberg *et al*, 2017).

To resolve the cellular specificity of *Zdbf2* expression, we used a previously described *LacZ* reporter *Zdbf2* gene-trap mouse line (Greenberg *et al*, 2017). X-gal staining of 2 week-old brain sections confirmed the expression of *Zdbf2* in various brain tissues and the high specificity in hypothalamic cells that belonged to the peri- and paraventricular nuclei in the anterior area (Figure 1B), and the arcuate, the dorsomedial and the ventromedial nuclei in the lateral hypothalamic area (Figure 1C). Analysis of publicly available single-cell RNA-sequencing (scRNA-seq) data further defined *Zdbf2* expression to be specific to neuronal cells and absent from non-neuronal cells of the hypothalamus (Figure S1B-C) (Chen et al., 2017). The three hypothalamic clusters where *Zdbf2* was the most highly expressed were both glutamatergic (Glu13 and Glu15) and GABAergic (GABA17) neurons from the arcuate and the periventricular hypothalamic regions (Figure S1B). Interestingly, these nuclei synthesize peptides that stimulate hormone production from the pituitary gland, or that control energy balance –food intake, energy expenditure and body temperature– by directly acting on the brain and/or more distal organs (Saper & Lowell, 2014).

**Figure 1:**
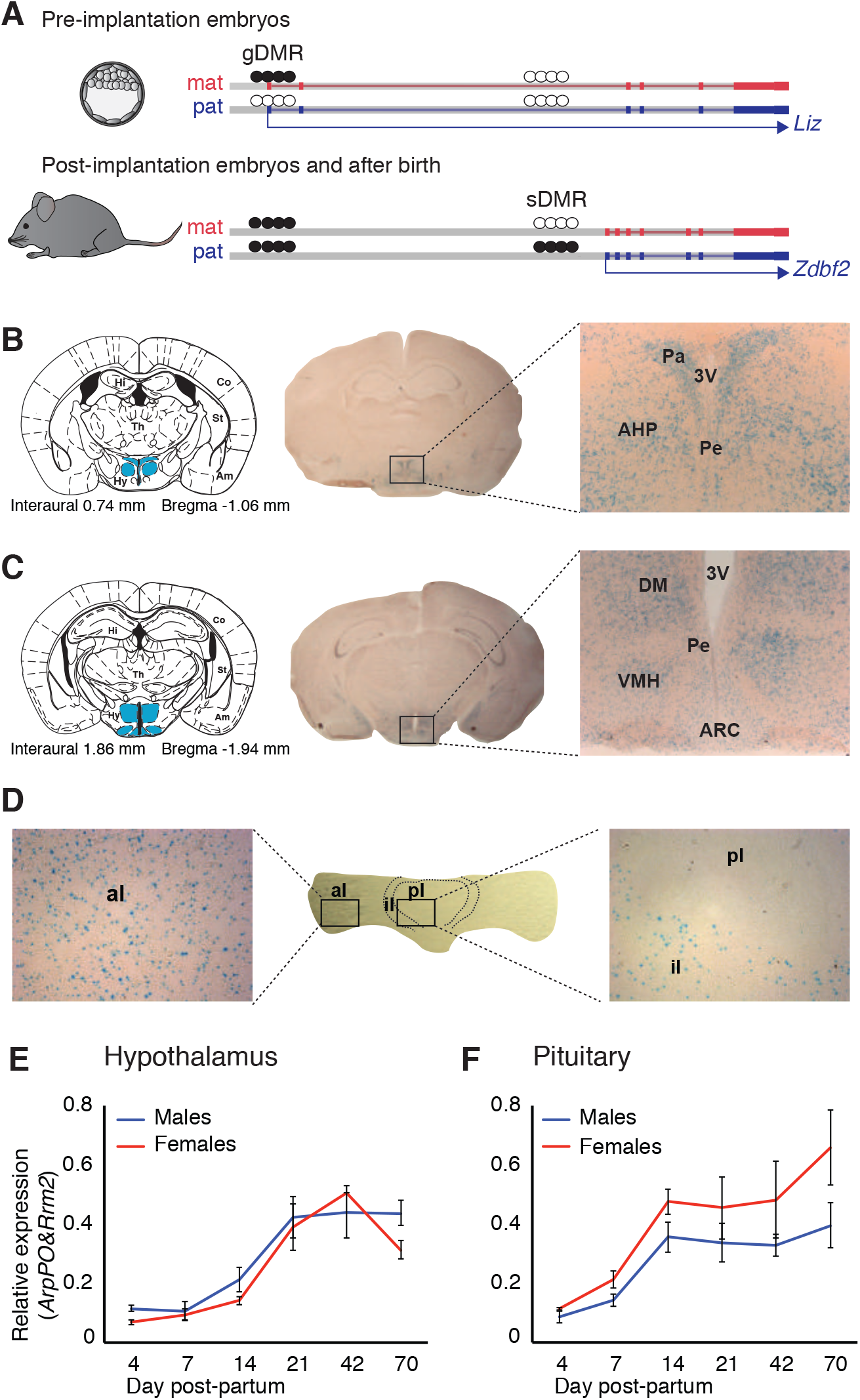
*Zdbf2* expression localizes preferentially in the neuro-endocrine cells of the hypothalamo-pituitary axis in juvenile animals. **(A)** Scheme of the *Liz/Zdbf2* locus regulation during mouse development. In the pre-implantation embryos, a maternally methylated gDMR allow the paternal-specific expression of the Long isoform of *Zdbf2* (*Liz*). *Liz* expression triggers, in cis, DNA methylation at the sDMR which is localized 8kb upstream of *Zdbf2* canonical promoter. In the post-implantation embryos and for the rest of the life, the imprint at the locus is controlled by the paternal methylation at the sDMR and this allow de-repression of *Zdbf2*, leading to its paternal-specific expression in the post-natal brain. (**B, C**) X-gal staining on brain coronal sections from 2-week-old *Zdbf2*-lacZ transgenic males. The coronal diagram from the Mouse Brain Atlas (left panel) localizes the sections in zone 40 (**B**) and zone 47 (**C**) in stereotaxic coordinates, with the hypothalamus indicated in blue. Whole brain coronal sections (middle panel) show specific staining in the hypothalamus, due to several positive hypothalamic nuclei (right panel, 20X magnificence). Hi, hippocampus; Co, cortex; Th, thalamus; Am, amygdala; St, striatum; Hy, hypothalamus; 3V, third ventricule; Pa, paraventricular hypothalamic nucleus; Pe, periventricular hypothalamic nucleus; AH, anterior hypothalamic area; DM, dorsomedial hypothalamic nucleus; VMH, ventromedial hypothalamic nucleus; Arc, arcuate hypothalamic nucleus. (**D**) X-gal staining on pituitary horizontal sections. The posterior lobe of the pituitary shows no X-gal staining (right panel), while staining is evenly distributed in the anterior lobe (left panel). pl, posterior lobe; il, intermediate lobe; al, anterior lobe. (**E, F**) *Zdbf2* expression measured by RT-qPCR in the hypothalamus (**E**) and the pituitary (**F**) from 4 to 70 days after birth. Data are shown as mean ± s.e.m. of n=3 C57Bl6/J mice.

*Zdbf2* being also expressed in the pituitary gland (Figure S1A), we hypothesized it could have a role in the hypothalamo-pituitary axis. When we exanimated the pituitary gland by X-gal staining, we found *Zdbf2* to be expressed in the anterior and intermediate lobes of the gland and almost undetectable signal in the posterior lobe (Figures 1D and S1D), a pattern that was confirmed from available scRNA-seq data (Figure S1E) (Cheung et al., 2018). The anterior and intermediate lobes that form the adenohypophysis are responsible for hormone production (Mollard *et al*, 2012). The anterior lobe secretes hormones from five different specialized hormone-producing cells under the control of hypothalamic inputs (growth hormone-GH, adrenocorticotropic hormone-ACTH, thyroid stimulating hormone-TSH, luteinizing hormone-LH and prolactin-PRL) and the intermediate lobe contains melanotrope cells (Kelberman *et al*, 2009). Altogether, the expression specificity of *Zdbf2* suggests a role in functions of the endocrine hypothalamo-pituitary axis and/or of the hypothalamus alone. Finally, we further found that steady-state levels of *Zdbf2* transcripts progressively rose in the hypothalamus and the pituitary gland after birth, reached their maximum at 2-3 weeks and then stabilized at later ages, in both males and females (Figures 1E and 1F). The expression of *Zdbf2* therefore mostly increases in juvenile pups prior to weaning.

### ZDBF2 positively regulates growth postnatally

In *Liz*-LOF mutants, *Zdbf2* failed to be activated and animals displayed postnatal body weight reduction (Greenberg *et al*, 2017). However, whether this was directly and only linked to *Zdbf2* deficiency was not resolved. To directly probe the biological role of ZDBF2, we therefore generated a mouse model of a genetic loss-of function of *Zdbf2*. More specifically, we engineered a ~700bp deletion of the entirety of exon 6 (Figure 2A) that is common to all annotated *Zdbf2* transcripts (Duffié *et al*, 2014). The *Zdbf2*-∆exon6 deletion induces a frame-shift predicted to generate a severely truncated protein (wild-type 2494aa versus 19aa mutant protein) that notably lacks the zinc finger motif. As *Zdbf2* is an imprinted gene with paternal-specific expression, the deletion should exhibit an effect upon paternal but not maternal transmission. For simplicity, heterozygous mutants with a paternally inherited *Zdbf2*-∆exon6 deletion are thus referred to as *Zdbf2*-KO thereafter.

**Figure 2:**
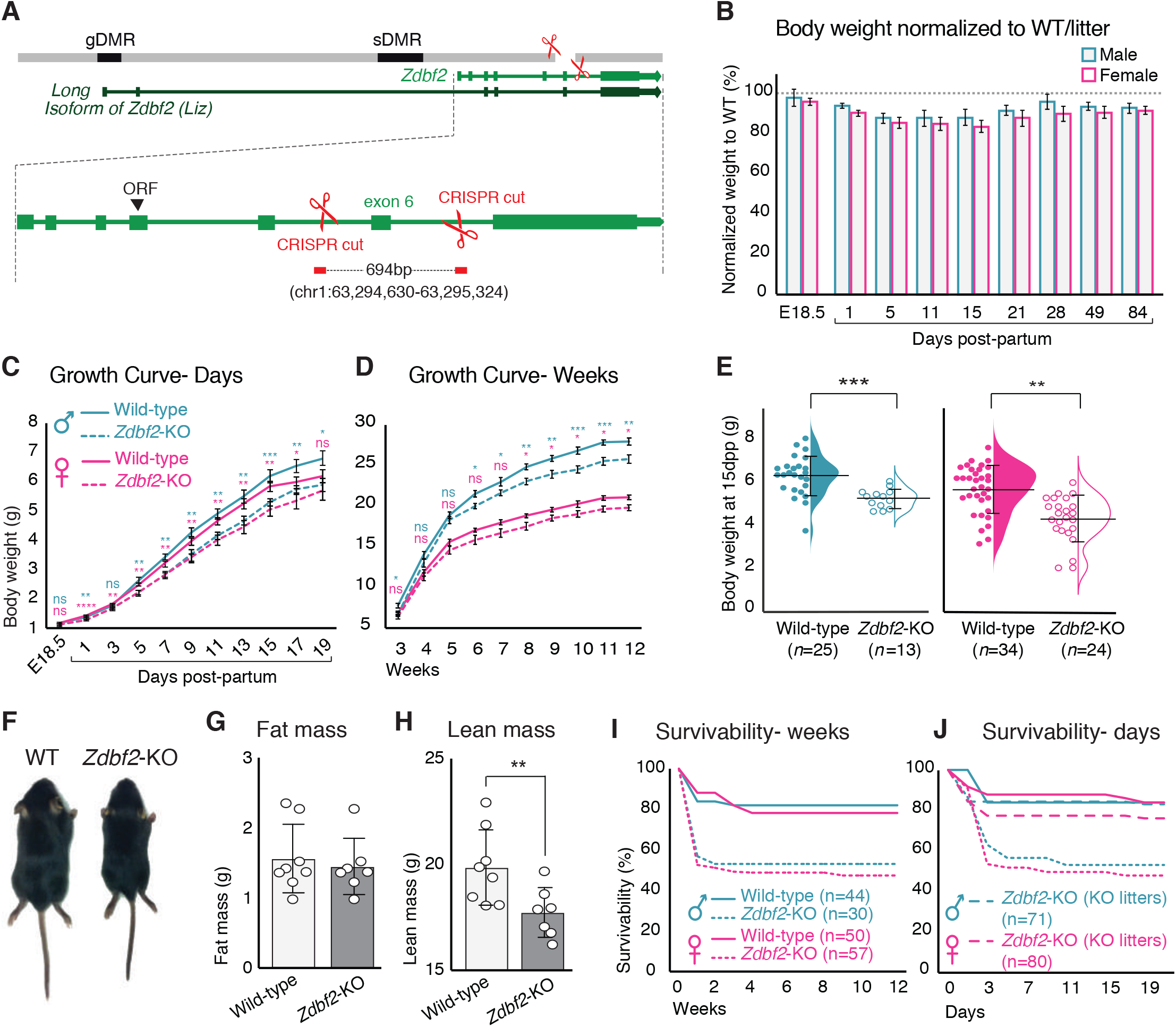
*Zdbf2*-KO mice exhibit growth reduction and partial post-natal lethality. (**A**) Graphical model of the *Zdbf2* deletion generated using two sgRNAs across exon 6. The two differentially methylated (DMR) regions of the locus are indicated (germline-gDMR, and somatic-sDMR), as well as the Long Isoform of *Zdbf2* (*Liz*). The ORF(Open Reading Frame) of *Zdbf2* starts in exon 4. Genomic coordinates of the deletion are indicated. (**B**) Body weight of *Zdbf2*-KO mice normalized to WT littermates (100%) followed from embryonic day E18.5 to 84 days post-partum. Data are shown as means ± s.e.m. from individuals from n= 27 litters. (**C, D**) Growth curve comparing the body weights of WT and *Zdbf2*-KO mice prior to weaning, from E18.5 to 19dpp (**C**) and over 3 months after birth (**D**). n=15–50 mice were analyzed per genotype, depending on age and sex. Statistical analyses were performed by a two-tailed, unpaired, nonparametric Mann Whitney t test. ***p ≤ 0.001,**p ≤ 0.01, *p≤ 0.05. (**E**) Half dot plot-half violin plot showing the weight distribution in 2 week-old males (left) and females (right) of WT and *Zdbf2*-KO genotypes. Statistical analyses were performed by a two-tailed, unpaired, nonparametric Mann Whitney t test. ***p ≤ 0.001,**p ≤ 0.01, *p≤ 0.05. (**F**) Representative photography of a smaller 2 week-old *Zdbf2*-KO male compared to a WT littermate. (**G-H**) Dual-energy X-ray absorptiometry (DXA) analysis showing the calculation of fat mass (**G**) and lean mass (**H**) in WT and *Zdbf2*-KO males at 7 weeks. Data are shown as means ± s.e.m. from n= 8 WT and n=7 *Zdbf2*-KO males. (**I**) Kaplan-Meier curves of the survivability from birth to 12 weeks of age comparing WT (plain lines) and *Zdbf2*-KO (dotted lines) littermates from WT x *Zdbf2* KO/WT backcrosses. Impaired survivability occurs specifically from the first day to 2 weeks of age. (**J**) Kaplan-Meier curves of the survivability from 1 to 21dpp comparing *Zdbf2*-KO pups generated from WT x *Zdbf2* KO/KO intercrosses, with *Zdbf2*-KO and their WT littermates generated from WT x *Zdbf2* KO/WT backcrosses. *Zdbf2*-KO pups are more prone to die only when they are in competition with WT littermates (small dotted lines), while *Zdbf2*-KO pups have a normal survivability when they are not with WT littermates (large dotted lines).

At birth, we found that *Zdbf2*-KO animals were present at expected sex and Mendelian ratios (Figures S2A and S2B). However, while exhibiting normal weight prior to birth (at embryonic day E18.5), *Zdbf2*-KO neonates of both sexes failed to thrive from the first day of postnatal life (day post-partum, dpp) (Figure 2B-D). Growth reduction was the most drastic prior to weaning, from 5 to 21dpp (Figure 2B): at 2 weeks of age, *Zdbf2*-KO mice were 20% lighter than their WT littermates (Figures 2B, 2E and 2F). Body weight reduction persisted through adulthood, with a tendency to minimize with age (Figure 2B). As predicted, when present on the maternal allele, the *Zdbf2*-∆exon 6 deletion had no discernable growth effect (Figures S2C and S2D).

The reduced body weight phenotype was fully penetrant (Figure S2E) and was not due to a developmental delay: *Zdbf2*-KO pups and their WT littermates synchronously acquired typical hallmarks of postnatal development (skin pigmentation, hair appearance and eye opening) (Figure S2F-I). The growth phenotype appeared to be systemic: it affected both the length and the weight of *Zdbf2*-KO animals (Figure S3A) and all organs uniformly (Figure S3B), such that body proportion was maintained. To gain insight into the origin of the body weight restriction, we performed dual-energy X-ray absorptiometry (DEXA) scan at 7 weeks of age to measure *in vivo* the volume fraction of the three dominant contributors to body composition: adipose, lean, and skeletal tissues (Chen et al., 2012). While we confirmed that adult *Zdbf2*-KO males exhibit an overall body weight reduction (Figure S3C), we did not observe difference in the composition of fat and bone tissues (Figures 2G and S3D-F). However, *Zdbf2*-KO mice had decreased lean mass (Figure 2H). Collectively, the above analyses demonstrate that the product of *Zdbf2* positively regulates body weight gain in juvenile mice.

In conclusion, *Zdbf2*-KO animals exhibit the same growth defect that we previously reported in *Liz*-LOF mice, with identical postnatal onset and severity (Greenberg *et al*, 2017). However, contrary to the *Liz*-LOF mice, the regulatory landscape of the paternal allele of *Zdbf2* was intact in *Zdbf2*-KO mice: DNA methylation of the sDMR located upstream of *Zdbf2* was normal (Figure S3G), and accordingly, *Zdbf2* transcription was activated (Figure S3H). This supports that *Liz* transcription in the early embryo –which is required for sDMR DNA methylation (Greenberg *et al*, 2017)— is not impacted in the *Zdbf2*-KO model. We therefore show that a genetic mutation of *Zdbf2* (*Zdbf2*-KO) and a failure to epigenetically program *Zdbf2* expression (*Liz*-LOF) are phenotypically indistinguishable. As *Liz*-LOF mice show a complete lack of *Zdbf2* expression (Greenberg *et al*, 2017), this incidentally supports that *Zdbf2*-KO mice carry a null *Zdbf2* allele. Unfortunately, we failed to specifically detect the mouse ZDBF2 protein with commercial or custom-made antibodies.

### *Zdbf2*-KO neonates fail to thrive or die prematurely within the first week of age

Growth restriction during the nursing period could dramatically impact the viability of *Zdbf2*-KO pups. Indeed, analysis of *Zdbf2*-KO cohorts revealed that although there was no Mendelian bias at 1dpp (Figure S2B), a strong bias was apparent at 20dpp: 75 WT and 44 *Zdbf2*-KO animals were weaned among *n*=27 litters, while a 50/50 ratio was expected. Post-natal survivability was the most strongly impaired within the first days after birth, with only 56% of *Zdbf2*-KO males and 52% of *Zdbf2*-KO females still alive at 3dpp, compared to 84 and 88% of WT survival at this age (Figure 2I and Supplemental Table 1A). Importantly, a partial postnatal lethality phenotype was also present in *Liz*-LOF mutants who are equally growth-restricted as a result of *Zdbf2* deficiency (Figure S3I), but did not occur when the mutation was transmitted from the silent maternal allele harboring either the *Zdbf2* deletion or the *Liz* deletion (Figure S3J and S3K and Supplemental Table 1C).

To determine whether partial lethality was due to a vital function of ZDBF2, *per se,* or to a competition with WT littermates, we monitored survivability in litters of only *Zdbf2*-KO pups generated from crosses between WT females x homozygous *Zdbf2*-KO/KO males. Litter sizes were similar than the ones sired by *Zdbf2*-KO/WT males. However, when placed in an environment without WT littermates, *Zdbf2*-KO pups gained normal survivability (Figure 2J and Supplemental Table 1B). Incidentally, this shows that *Zdbf2*-KO pups have the mechanical ability to suckle milk. More likely, the impaired survival of *Zdbf2*-KO neonates is a consequence of intra-litter competition with healthier WT pups. Overall, our detailed study of the *Zdbf2*-KO phenotype reveals that smaller *Zdbf2*-KO neonates have reduced fitness compared to their WT littermates. In absence of ZDBF2, after a week of life, half of juvenile pups did not survive and the other half failed to thrive.

### Postnatal body weight control is highly sensitive to *Zdbf2* dosage

The physiological effects of imprinted genes are intrinsically sensitive to both decreases and increases in the expression of imprinted genes. We therefore went on assessing how postnatal body weight gain responded to varying doses of the imprinted *Zdbf2* gene. We first analyzed the growth phenotype resulting from partial reduction in *Zdbf2* dosage, as compared to a total loss of *Zdbf2* in *Zdbf2*-KO mice. For this, we used the *LacZ* reporter *Zdbf2* gene-trap mouse line (Greenberg *et al*, 2017), in which the insertion of the *LacZ* cassette downstream of exon 5 does not fully abrogate the production of full-length *Zdbf2* transcripts (Figure S4A). Upon paternal inheritance of this allele, animals still express 50% of WT full-length *Zdbf2* mRNA levels in the hypothalamus and the pituitary gland (Figure S4B and S4C). Interestingly, these mice displayed significant weight reduction compared to their WT littermates (Figure S4D), but less severe than *Zdbf2*-KO mice (Figure 2B-D). On average, we observed a 5 to 10% weight reduction from 7 to 21dpp, which is incidentally half the growth reduction observed in *Zdbf2*-KO mice for the same age range.

We then assessed the consequences of increased *Zdbf2* dosage, with the hypothesis that this would lead to excessive body weight gain after birth, as opposed to reduced *Zdbf2*. For this, we took advantage of a *Zdbf2* gain-of-function (GOF) line that we serendipitously obtained when generating *Liz* mutant mice (Greenberg *et al*, 2017) (see detailed description in Supplemental information). The *Zdbf2*-GOF lines carry a ~ 900bp deletion upstream of *Liz* exon 1, which leaves some part of exon 1 intact and induces an epigenetic “paternalization” of the maternal allele of the locus (Figure S4E and S4F). While sDMR methylation and *Zdbf2* activation exclusively occur on the paternal allele during normal development, animals that maternally inherit this partial deletion also acquire sDMR methylation on the maternal allele (Figures 3A and S4F-H) and activate *Zdbf2* from the maternal allele, although slightly less than from the WT paternal allele (Figure 3B). As a consequence, *Zdbf2*-GOF animals exhibit bi-allelic *Zdbf2* expression, with a net 1.7-fold increase of *Zdbf2* levels in postnatal hypothalamus and pituitary gland, as compared to WT littermates that express *Zdbf2* mono-allelically, from the paternal allele only (Figure 3C).

**Figure 3:**
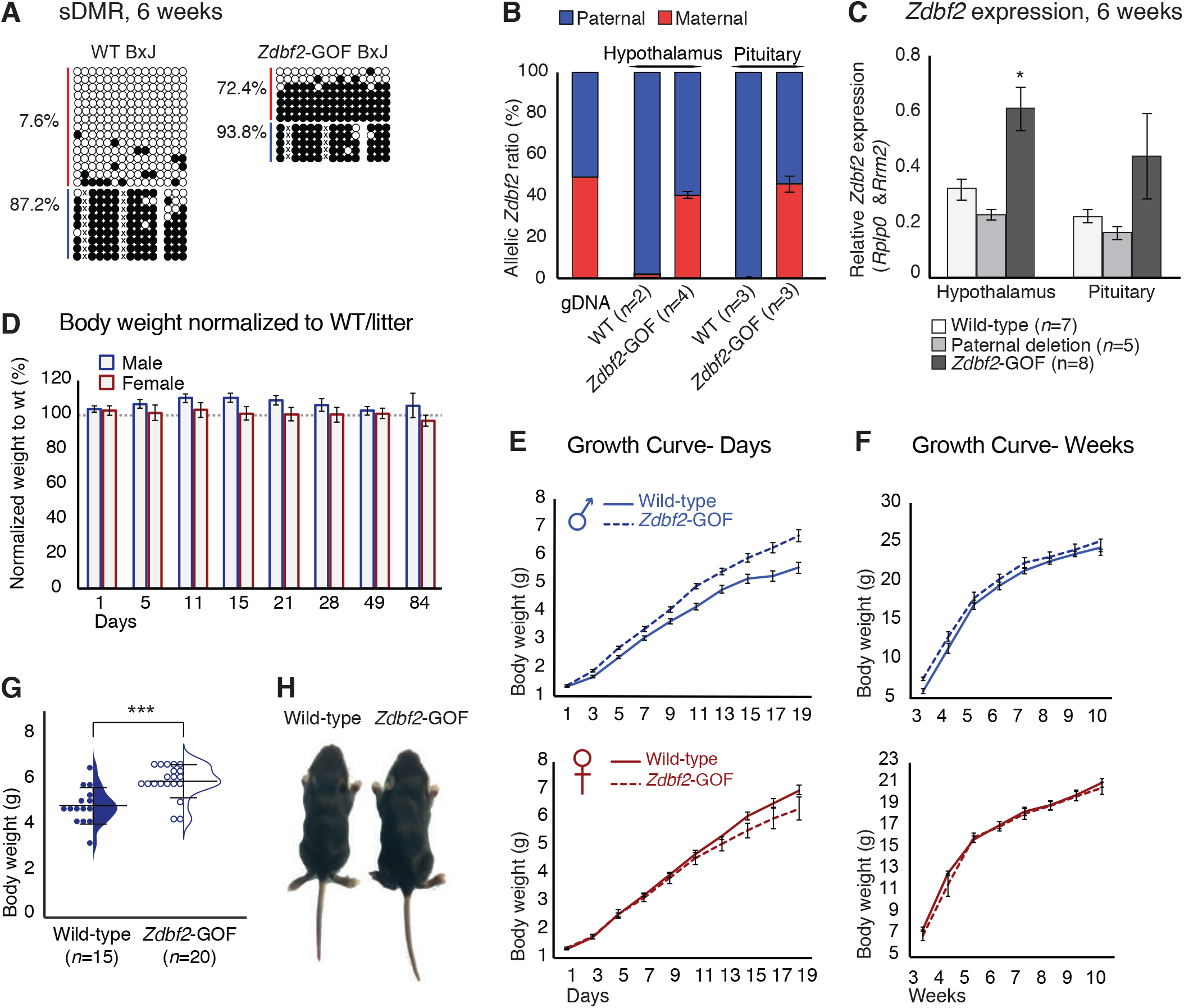
*Zdbf2* influences postnatal growth in a dose-dependent manner. (**A**) Bisulfite cloning and sequencing showing CpG methylation levels at the sDMR locus of hypothalamus DNA from 6-week-old hybrid WT (left) and *Zdbf2*-GOF (right) mice (*Zdbf2*-GOF +/-x JF1 cross). Red, maternal alleles; blue, paternal alleles. cross, informative JF1 SNP. (**B**) Allelic expression of *Zdbf2* in hypothalamus and pituitary gland from 3-week-old mice, measured by RT-pyrosequencing. Genomic DNA extracted from a C57Bl/6 x JF1 hybrid cross was used as a control for pyrosequencing bias. (**C**) RT-qPCR measurement reveals a ~ 1.7 fold-increase of *Zdbf2* expression in the hypothalamus and pituitary gland of 3 week-old mice with a maternal transmission of the deletion. Expression of *Zdbf2* in mice carrying the deletion on the paternal allele is similar to WT. (**D**) Normalized body growth of *Zdbf2*-GOF mice to their WT littermates (100%) followed at different ages (1 to 84 days) from n=14 litters. The overgrowth is seen specifically in males, from 5 to 28 days. (**E, F**) Growth curves of female and male mice, comparing the body weights of WT and *Zdbf2*-GOF, through the 3 first weeks of life (D) and through 10 weeks (E). n=10–30 mice were analyzed per genotype, depending on age and sex. (**G**) Half dot-half violin plots showing the weight distribution at 2 weeks of age between WT and *Zdbf2*-GOF males. Data are shown as means ± s.e.m. from n individuals. Statistical analyses were performed by a two-tailed, unpaired, nonparametric Mann Whitney t test. *** p ≤ 0.005. (**H**) Representative photography of a bigger *Zdbf2*-GOF male as compared to a WT littermate at 2 weeks.

Importantly, we found that increased *Zdbf2* dosage was indeed impactful for postnatal body weight: *Zdbf2*-GOF animals were consistently overweight (Figure 3D-F), and their viability was normal (Figure S5A-C). Although *Zdbf2* was overexpressed in both *Zdbf2*-GOF males and females, the overgrowth phenotype was male-specific, which may imply stimulation of the phenotype by sex hormones, directly or indirectly. Similar to growth restriction observed in *Zdbf2*-KO mutant mice, increased body weight of *Zdbf2*-GOF males was more pronounced during the nursing period (Figure 3D, G and H) and affected all organs uniformly (Figure S5D). From 10 to 20dpp, *Zdbf2*-GOF juvenile males were 10% heavier when normalized to the average WT siblings from the same litter (Figure 3D). At 2 weeks of age, *Zdbf2*-GOF males were visibly bigger than their WT littermates (Figure 3G and 3H), while females displayed standard normal distribution (Figure S5E). Comparatively, progenies from a paternal transmission of the deletion showed a growth progression similar to their WT littermates (Figure S5F and S5G). Overall, these data illustrate the importance of *Zdbf2* for the regulation of postnatal body weight gain at the onset of postnatal life. We therefore demonstrate that the *Zdbf2* imprinted gene encodes a genuine positive regulator of postnatal body weight gain in mice, with highly attuned dose-sensitive effects.

### *Zdbf2* regulates post-natal growth in a parent-of-origin independent manner

Imprinted genes with a paternal expression generally tend to enhance prenatal and postnatal growth (Haig, 2000). Consistently, we found that the paternally expressed *Zdbf2* gene promotes postnatal weight gain. Having determined the dosage effect of *Zdbf2*, we next wondered what role plays the paternal origin of *Zdbf2* expression on postnatal body weight. To invert *Zdbf2* parental expression, we intercrossed the *Liz*-LOF and *Zdbf2*-GOF lines, which consist in a maternalization of the paternal allele and a paternalization of the maternal allele of *Zdbf2*, respectively. By crossing an heterozygous *Zdbf2*-GOF female with an heterozygous *Liz*-LOF male (Figure 4A, left panel), four genotypes can segregate in the progeny, with littermates displaying various dosage and parent-of-origin expression of *Zdbf2* (Figure 4A): *1)* wild-type animals with one dose of paternal *Zdbf2* expression, *2)* animals with a paternal *Liz*-LOF allele and lack of *Zdbf2* expression (equivalent to a *Zdbf2*-loss-of-function, *Zdbf2*-LOF), *3)* animals with a maternal *Zdbf2*-GOF allele and biallelic *Zdbf2* expression, and finally, *4)* animals with combined maternal *Zdbf2*-GOF and paternal *Zdbf2-*LOF alleles and potentially, mono-allelic, reverted maternal expression of *Zdbf2* (Figure 4A, right panel).

**Figure 4:**
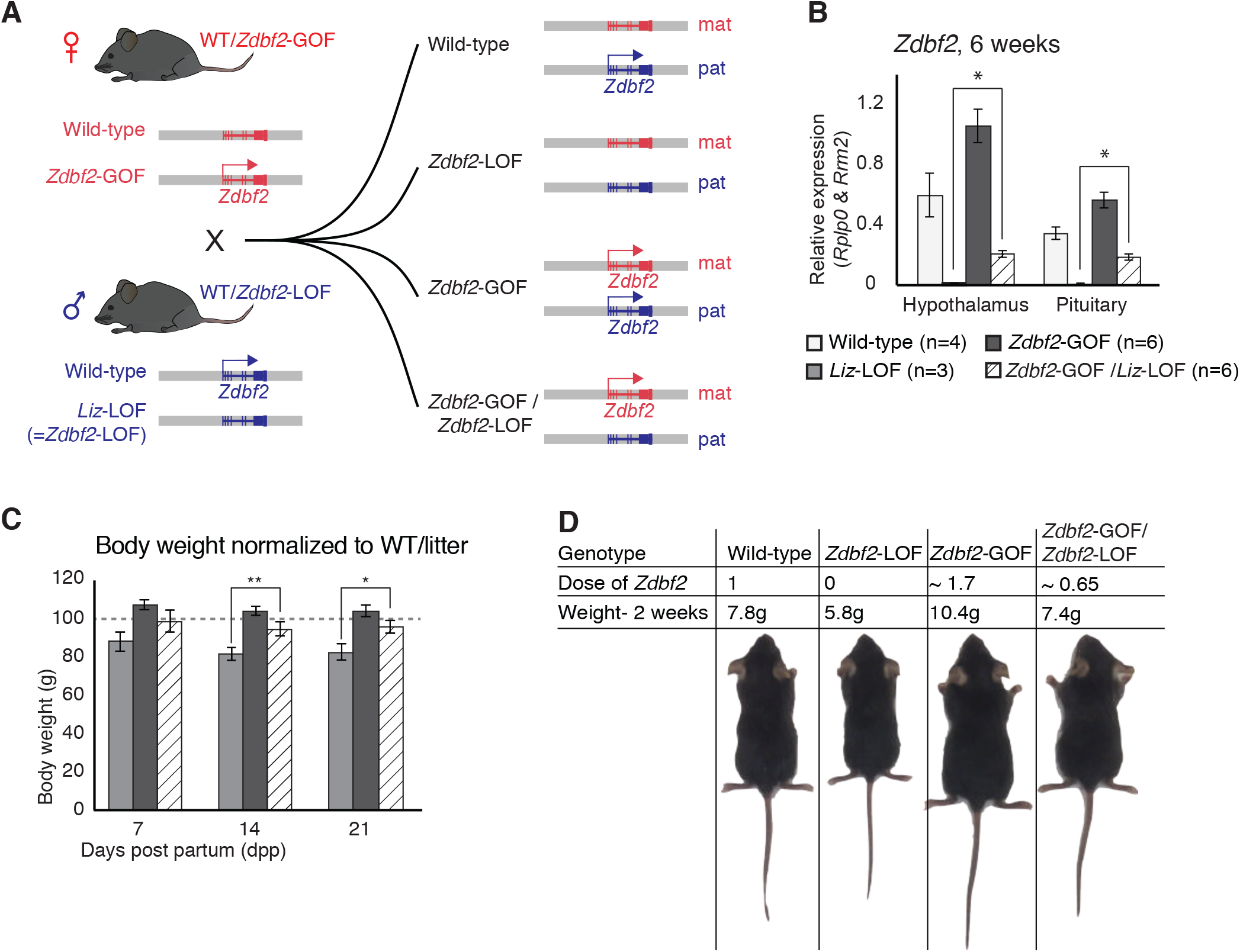
*Zdbf2* influences postnatal growth in a parent-of-origin-independent manner. (**A**) Scheme of the cross made to obtain embryos with an inversion of the parental origin of *Zdbf2* expression. *Zdbf2*-GOF heterozygote females were crossed with heterozygote males for the *Liz*-LOF deletion which we demonstrated as equivalent to a *Zdbf2*-LOF allele- (left) to obtain one quarter of embryos expressing one dose of *Zdbf2* from the maternal allele (right, bottom). (**B**) *Zdbf2* expression in the hypothalamus and the pituitary gland is shown in males for each of the four possible genotypes. The level of *Zdbf2* in *Zdbf2*-GOF /*Zdbf2-*LOF mice almost completely rescues the defect seen in *Zdbf2*-LOF and *Zdbf2*-GOF mutants. Data are shown as means ± s.e.m. from n individuals. Statistical analyses were performed by a two-tailed, unpaired, nonparametric Mann Whitney t test. * p ≤ 0.05. (**C**) Normalized body growth of *Zdbf2*-LOF, *Zdbf2*-GOF and *Zdbf2*-GOF /*Zdbf2*-LOF males to their WT littermates (100%) followed at 7, 14 and 21 days after birth. *Zdbf2*-GOF /*Zdbf2*-LOF adult mice exhibit a body weight similar to the WT showing a partial rescue of the growth reduction and overgrowth phenotype due to respectively the lack of *Zdbf2* and the gain of *Zdbf2* expression in the brain. Data are shown as means ± s.e.m. from individuals from n= 17 litters. Statistical analyses were performed by a two-tailed, unpaired, nonparametric Mann Whitney t test. *p ≤ 0.05; **p ≤ 0.01. (**D**) Representative photography of four males littermates from a *Zdbf2*-GOF x *Zdbf2*-LOF cross (as shown in A) at 2 weeks of age. For each animal, genotype, dose of *Zdbf2* expression and weight are indicated.

The uniform strain background of the two lines did not allow us to use strain-specific sequence polymorphisms to distinguish the parental origin of *Zdbf2* regulation in the compound *Zdbf2*-GOF/*Zdbf2*-LOF animals. However, we found that compared to single *Zdbf2*-LOF mutants, the presence of the maternal *Zdbf2*-GOF allele restored DNA methylation levels at the sDMR locus in all tissues of *Zdbf2*-GOF/*Zdbf2*-LOF mutants, with an average of 43.5% CpG methylation compared to the expected 50% in WT (Duffié *et al*, 2014) (Figure S5H). By RT-qPCR, *Zdbf2* mRNA levels were also increased in the hypothalamus and pituitary gland of *Zdbf2*-GOF/*Zdbf2*-LOF animals compared to single *Zdbf2*-LOF animals (Figure 4B), which strongly suggests that expression comes from the maternal *Zdbf2*-GOF allele. Accordingly, *Zdbf2* expression level in *Zdbf2*-GOF/*Zdbf2*-LOF animals was on average 0.65-fold the one of WT animals, which is congruent with the partial paternalization of the maternal *Zdbf2*-GOF allele we reported (Figures 4B and S4I). Most importantly, restoration of *Zdbf2* expression by the maternal *Zdbf2*-GOF allele -even though incomplete- was sufficient to rescue the postnatal body weight phenotype in compound *Zdbf2*-GOF/*Zdbf2*-LOF males compared to their single *Zdbf2*-LOF brothers (Figure 4C and 4D). From a body weight reduction of 20% reported in *Liz*-LOF or *Zdbf2*-KO animals at the same age, it was attenuated to only 4% in *Zdbf2*-GOF/*Zdbf2*-LOF animals (Figure 4C), showing that maternal *Zdbf2* expression is as functional as paternal *Zdbf2* expression. Altogether, these results crystallize the importance of *Zdbf2* dosage in regulating postnatal body weight and most importantly, demonstrate that the dose but not the parental origin matters for *Zdbf2* function.

### *Zdbf2*-KO growth phenotype is linked to decreased IGF-1 in the context of normal development of the hypothalamo-pituitary axis

Having demonstrated the growth promoting effect of *Zdbf2*, we next tackle the question of how it influences newborns weight and survival. As *Zdbf2* is expressed in the neuroendocrine cells of the hypothalamo-pituitary axis, the phenotype of *Zdbf2*-KO animals may lay in a defect in producing growth-stimulating pituitary hormones. When we assessed the development and functionality of the hypothalamus and pituitary gland prior to birth, we did not detect any morphological nor histological defects in *Zdbf2*-KO embryos (Figure S6A-C). Normal expression of major transcriptional regulators confirmed proper cell lineage differentiation in the *Zdbf2*-KO developing pituitary (Figure S6B) (Raetzman *et al*, 2002; Rizzoti, 2015; Kelberman *et al*, 2009). Immunohistochemistry at E18.5 further indicated that *Zdbf2*-KO pituitary cells acquire normal competency for producing hormones (Figure S6D). Similarly, hypothalamic peptides were expressed in comparable levels in *Zdbf2*-KO and WT embryos, as assessed by *in situ* hybridization analysis (Figure S6E and S6F) (Biran *et al*, 2015). In sum, the embryonic hypothalamo-pituitary axis develops normally in the absence of *Zdbf2*. Our data implies that the postnatal growth phenotype does not result from impaired establishment or programming of this axis during embryogenesis.

We then went on to analyze the functionality of the hypothalamo-pituitary axis in producing hormones after birth. Again, GH, ACTH, TSH, LH and PRL all appeared to be normally expressed in the pituitary glands of *Zdbf2*-KO juvenile animals (immunohistochemistry at 15dpp) (Figure 5A). Although we cannot exclude subtle dysfunctionalities, our results suggest that *Zdbf2* deficiency does not compromise pituitary hormone production. Hormone release was also functional, as we measured normal plasmatic GH levels in 5 and 15dpp *Zdbf2*-KO animals (Figure 5B). We therefore excluded an involvement of GH in the *Zdbf2*-KO growth phenotype.

**Figure 5:**
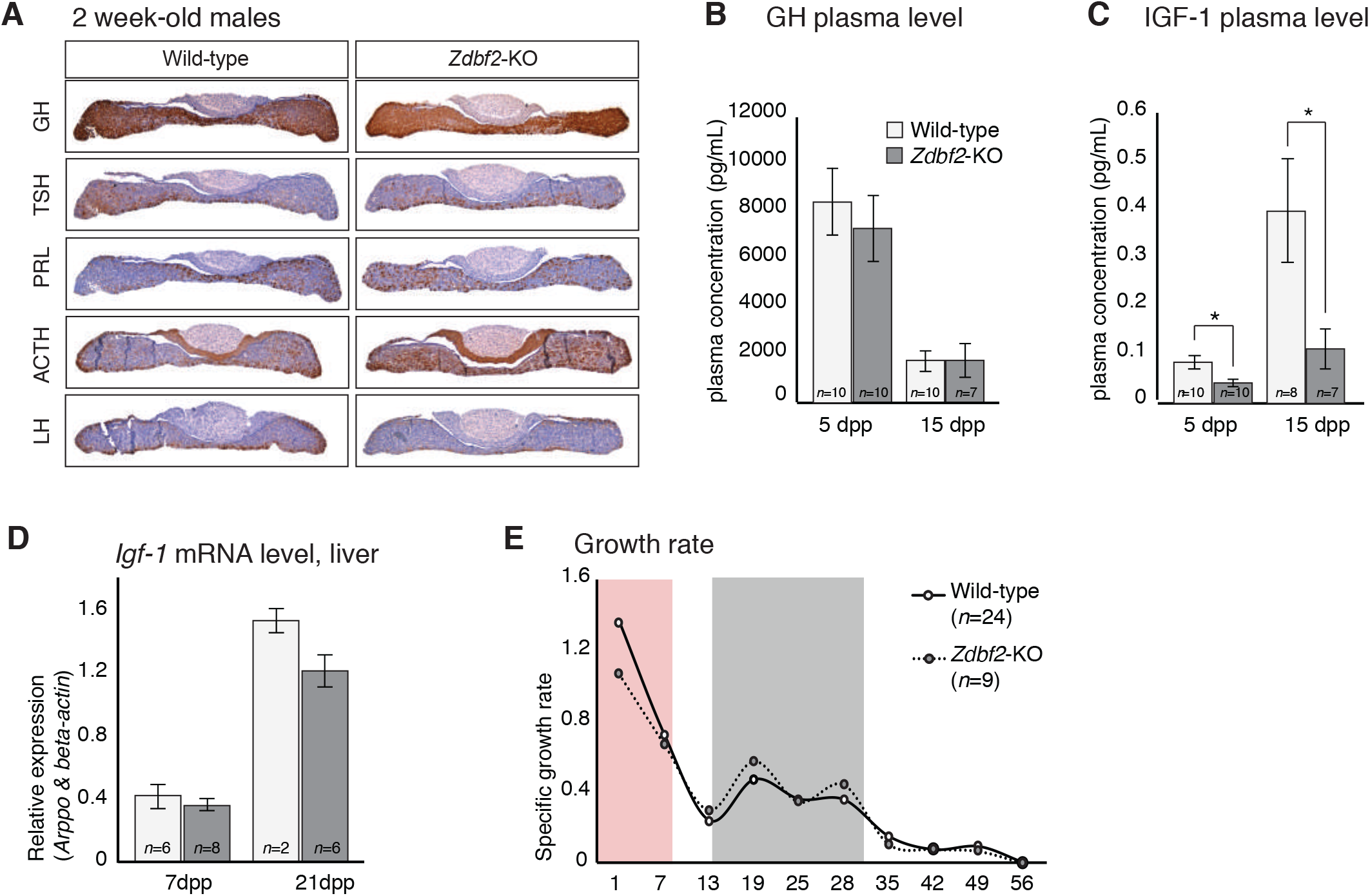
*Zdbf2*-KO phenotype is linked to defective IGF-1 signaling immediately after birth. (**A**) Pituitary hormone production is globally normal in *Zdbf2*-KO mice, as assessed by immunohistochemistry on 15dpp pituitary sections. (**B, C**) Circulating levels of GH (**B**) and IGF-1 (**C**) in the plasma of WT and *Zdbf2*-KO mice at 5 and 15dpp. Data are shown as means ± s.e.m. from n replicates. Statistical analyses were performed by a two-tailed, unpaired, nonparametric Mann Whitney t test.*p ≤ 0.05. (**D**) RT-qPCR from post-natal liver measuring the level of *Igf-1* mRNAs between WT and *Zdbf2*-KO mice at 7 and 21dpp. Data are shown as means ±s.e.m from n WT and *Zdbf2*-KO animals. (**E**) Specific growth rates calculated from body weight of WT and *Zdbf2*-KO males from 1 to 56dpp using the following equation: [(weight t2 - weight t1)/ weight t1]. Pink area: reduced growth rate in *Zdbf2*-KO pups compared to WT littermates; Grey area: growth rate of *Zdbf2*-KO mice exceeds the WT growth rate.

However, *Zdbf2*-KO pups showed reduced plasma circulating levels of insulin growth factor 1 (IGF-1) –a main marker of postnatal growth– which reached only 50% and 20% of WT level at 5dpp and 15dpp, respectively (Figure 5C). In adult mice, the liver is the main site of production of IGF-1, following transcriptional activation under the control of circulating GH (Savage, 2013). In contrast, extrahepatic production of IGF-1 during embryonic and early postnatal life is mostly GH-insensitive (Lupu *et al*, 2001; Kaplan & Cohen, 2007). As mentioned above, decreased IGF-1 levels in juvenile *Zdbf2*-KO animals indeed occurs in the context of normal GH input. Moreover, we measured normal *Igf1* mRNA levels by RT-qPCR in the liver of juvenile *Zdbf2*-KO animals, further illustrating that decreased IGF-1 levels are not a result of altered GH pathway (Figure 5D).

The association of low levels of circulating IGF-1 with normal GH secretion prompted us to evaluate more thoroughly the *Zdbf2*-KO growth phenotype. Mouse models of GH deficiency show growth retardation only from 10dpp onwards, while deficiency in IGF-1 affects growth earlier during postnatal development (Lupu *et al*, 2001). When we calculated the growth rate from day 1 to 8 weeks of age, we revealed two distinct phases: *(1)* from 1 to 7dpp, the *Zdbf2* mutant growth rate was 22% lower compared to WT littermates and *(2)* from 15 to 35dpp, the mutant exceeded the WT growth rate (Figure 5E) (catch up phase), showing that the growth defect is exquisitely restrained to the first days of life. More specifically, at the day of birth (1dpp), *Zdbf2*-KO pups were smaller than their WT littermates by 9% (WT 1.41 ± 0.015g, *n*=99; *Zdbf2*-KO 1.29 ± 0.013, *n*=95). After 1 week of post-natal life, the mutant growth restriction reached 18% (WT 3.4 ± 0.13g, *n*=37; *Zdbf2*-KO 2.8 ± 0.11, *n*=17), indicating a failure to thrive prior to 10dpp. Then, at 6 weeks of age, *Zdbf2*-KO animals were smaller than WT by only 8%, as a result of enhanced post-pubertal growth spurt (WT 21.7 ± 0.04g, *n*=34; *Zdbf2*-KO 20.05 ± 0.3, *n*=15). We therefore concluded that the *Zdbf2*-KO phenotype is similar to defective IGF-1 signaling immediately after birth.

Overall, we revealed that the body weight restriction of *Zdbf2*-KO juveniles is associated with a GH-independent decrease of IGF-1 during the first days of postnatal life, in the context of an overall normal development and functionality of the hypothalamo-pituitary axis.

### *Zdbf2*-KO neonates are undernourished and do not properly activate hypothalamic feeding circuits

Having determined that *Zdbf2*-KO animals exhibit an *Igf-1*-like phenotype, we attempted to define the origin of the GH-independent reduction of IGF-1 levels. Undernutrition is a well-known cause of IGF-1 level reduction, and also of postnatal lethality (Thissen *et al*, 1994), which we observed in *Zdbf2*-KO pups. To investigate the nutritional status of *Zdbf2*-KO neonates, we weighed stomachs at 3dpp, as a measure of milk intake. *Zdbf2*-KO pups exhibited a significant reduction in stomach weight relative to body mass as compared to their WT littermates (Figures 6A and S7A), suggesting that these pups suffer milk deprivation. Comparatively, other organs showed normal relative weight at this age, at the exception of the interscapular brown adipose tissue (BAT), which was also reduced in *Zdbf2*-KO neonates (Figures 6B and S7A). BAT-mediated thermogenesis regulates body heat during the first days after birth (Cannon & Nedergaard, 2004) and improper BAT function can lead to early postnatal death (Charalambous *et al*, 2012). However, despite being smaller, the BAT of *Zdbf2*-KO neonates appeared otherwise functional, showing normal lipid droplet enrichment on histological sections (data not shown) and proper expression of major markers of BAT thermogenic ability (Figure S7B). The BAT size reduction may therefore not reflect altered BAT ontogeny *per se,* but rather the nutritional deprivation of *Zdbf2*-KO neonates.

**Figure 6:**
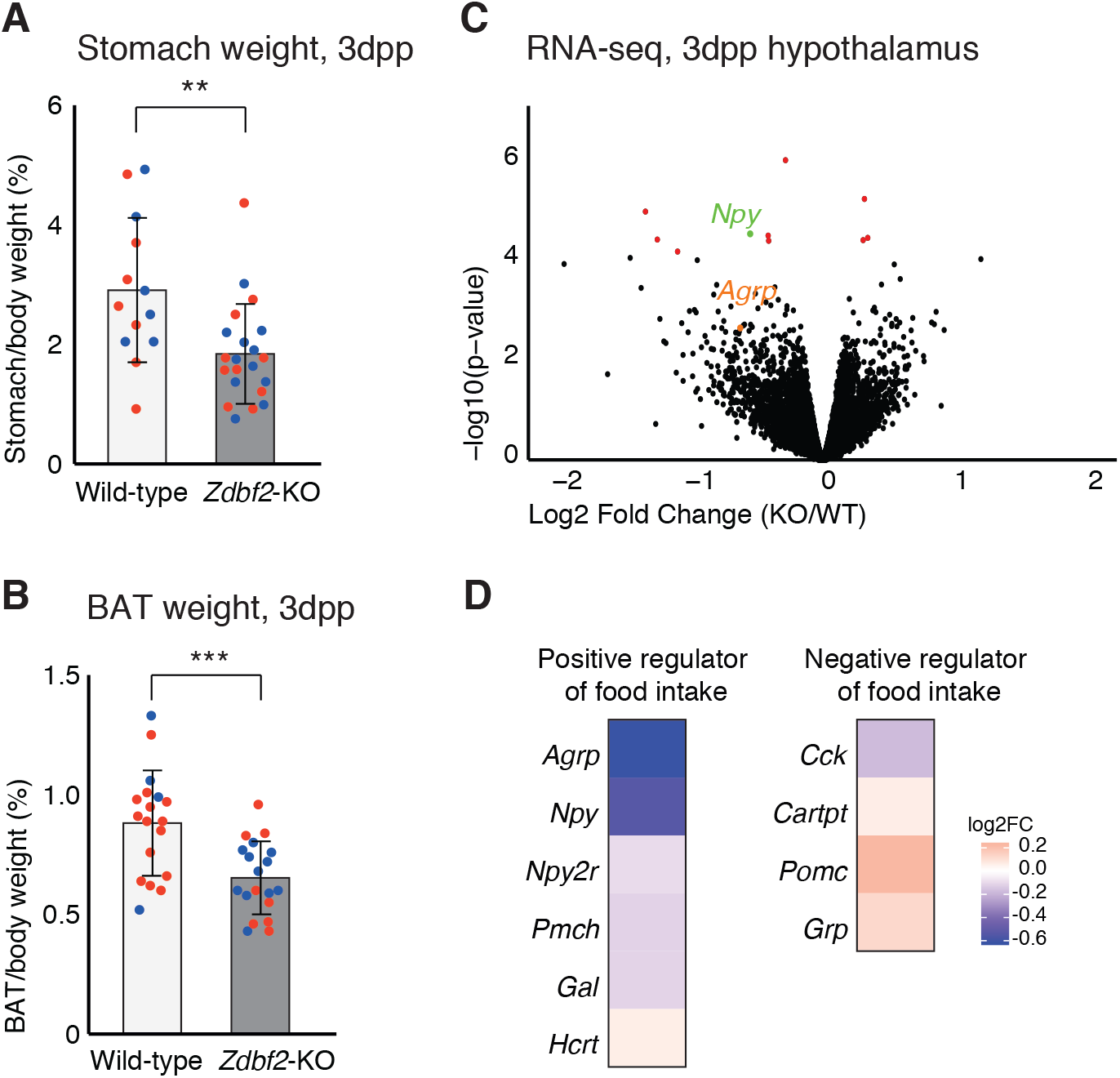
*Zdbf2*-KO neonates display a feeding defect. (**A, B**) Stomach (**A**) and brown adipocyte tissue (BAT) (**B**) weight normalized to the body weight for WT and *Zdbf2*-KO at 3dpp. Red dots: females; blue dots: males. Data are shown as means ± s.e.m. from n replicates. Statistical analyses were performed by a two-tailed, unpaired, nonparametric Mann Whitney t test.** p≤ 0.01, ***p ≤ 0.005. (**C**) Volcano plot representation of RNA-seq of 3dpp hypothalamus of *Zdbf2*-KO versus WT littermates. Red dots: differentially expressed genes with a threshold of FDR<10%. Npy (FDR 6%) is highlighted in green and Agrp in orange. (**D**) Heatmap showing the log2 fold change of genes encoding hypothalamic regulators of food intake (RNA-seq data from C).

Because the mothers of *Zdbf2*-KO pups are of WT background, the undernutrition phenotype is unlikely due to defective maternal milk supply. Restoration of viability when placed in presence of KO-only littermates (Figure 2K) further indicated that *Zdbf2*-KO pups are competent for suckling. Defective nutrition may therefore rather result from altered feeding motivation of the *Zdbf2*-KO neonates, specifically during the early nursing period. To test this hypothesis, we performed RNA-seq analysis of dissected hypothalami at 3dpp, when the growth phenotype is the most acute (Figure 6C). Only 11 genes were significantly misexpressed in the *Zdbf2*-KO pups relative to their WT littermates (FDR 10%), with a majority being downregulated genes, including *Npy*, which encodes the Neuropeptide Y. In line with our hypothesis, NPY is secreted from neurons of the hypothalamic arcuate nucleus, where *Zdbf2* is expressed and stimulates food intake and promotes gain weight (Mercer *et al*, 2011). Little is known about the determinants of food intake in neonates (Muscatelli & Bouret, 2018), nonetheless, this prompted us to examine the expression levels of other hypothalamic modulators known to influence feeding behavior in adults. Interestingly, *Zdbf2*-KO pups showed a trend towards downregulation of the Agouti-related peptide (*Agrp*)-encoding gene (Figure 6D), which is co-expressed with *Npy* in a sub-population of hypothalamic neurons and convergently stimulate appetite and food seeking (Gropp *et al*, 2005). It is noteworthy that none of the misregulated genes –including positive regulators of feeding– that we observed at 3dpp were continuously misregulated in the hypothalamus at 10dpp, when *Zdbf2*-KO pups enter the catch-up phase (Figure S7C and S7D).

Together, these results provide key insights into the origin of the *Zdbf2* mutant phenotype: in absence of *Zdbf2,* the hypothalamic circuit of genes that stimulates food intake may not be properly activated at birth. This is associated with reduced milk intake, reduced body weight gain and suboptimal viability of *Zdbf2*-KO neonates when placed in presence of healthier littermates.

## DISCUSSION

The hypothalamus is increasingly regarded as a major site for the action of imprinted genes on postnatal growth, feeding behavior and metabolism (Ivanova & Kelsey, 2011). Here, we found evidence that the imprinted *Zdbf2* gene stimulates hypothalamic feeding circuits, right after birth. In its absence, neonates do not suckle enough milk, and suffer from undernutrition. This leads to decreased IGF-1 signaling within the first week of life, reduced body weight gain and more dramatically, lethality of half of the *Zdbf2*-KO pups when they are in presence of healthier WT littermates. Our work therefore highlights that *Zdbf2* is necessary to thrive and survive after birth, by allowing newborns to adapt to postnatal feeding.

By relying on a unique collection of mouse models of total loss of function (*Zdbf2-KO* and *Liz-LOF*), partial loss of function (*Zdbf2*-lacZ), normal function (*Zdbf2*-WT) and gain of function (*Zdbf2*-GOF), we observed that postnatal body growth is exquisitely sensitive to the quantity of *Zdbf2* produced in the hypothalamo-pituitary axis. Incidentally, *Zdbf2* meets the criteria of a *bona fide* growth-promoting gene: growth is reduced upon decreased dosage and oppositely enhanced upon increased dosage of *Zdbf2* (Efstratiadis, 1998). Consistent with previous results for the imprinted *Cdkn1c* gene (Andrews *et al*, 2007), growth reduction was more pronounced than overgrowth upon changes in *Zdbf2* expression (18% decrease and 10% increase at 10dpp) and specific to males. This reflects that, with standard diet, overgrowth is a much less frequent response than growth reduction and affects animals with favorable physiology only, potentially explaining male specificity.

With a few exceptions, most studies have only addressed the phenotypic effects of reducing the dose of an imprinted gene, but not what results from overexpressing this same gene (Tucci *et al*, 2019). Knocking out an imprinted gene provides valuable insights about its physiological function, however, it does not address the evolutionary significance of imprinting of that gene, *i.e.* reduction to mono-allelic expression. With the *Zdbf2*-GOF model, we were able to evaluate the consequences of a loss of *Zdbf2* imprinting: bi-allelic and increased *Zdbf2* expression exacerbates postnatal body weight gain. However, despite being slightly bigger, viability, fertility or longevity appeared overall normal in *Zdbf2*-GOF animals. The evolutionary importance of *Zdbf2* imprinting is therefore not immediately obvious, at least under unchallenged conditions. Finally, even fewer studies have addressed the importance of parent-of-origin expression for imprinted gene functions (Drake *et al*, 2009; Leighton *et al*, 1995). By intercrossing models of *Zdbf2* loss and gain of function, we could enforce *Zdbf2* expression from the maternal allele while inactivating the normally expressed paternal allele. Restoration of body weight demonstrated the functionality of maternal *Zdbf2* expression, validating that *Zdbf2* functions on postnatal body homeostasis in a dose-dependent but parent-of-origin-independent manner. Although not necessarily surprising, this demonstration is conceptually important for understanding the evolution and *raison d’être* of genomic imprinting. The divergent DNA methylation patterns that are established in the oocyte and the spermatozoa provide opportunities to evolve mono-allelic regulation of expression, but the parental information is not essential *per se*.

Although *Zdbf2* is expressed across the hypothalamo-pituitary axis, we could not find evidence of abnormal development or function of the pituitary gland that could explain the *Zdbf2*-KO growth phenotype. Notably, GH production and release was normal. Additionally, the *Zdbf2* growth reduction diverges from GH-related dwarfism: *Gh*-deficient mice grow normally until 10dpp, after which only they exhibit general growth impairment with reduced levels of circulating IGF-1 (Voss & Rosenfeld, 1992; Lupu *et al*, 2001). In *Zdbf2*-KO mutants, the growth defect is apparent as soon as 1dpp and we measured decreased IGF-1 at 5 dpp. In *Igf-1* null mice, birthweight is approximately 60% of normal weight and some mutants die within the first hours after birth (Liu *et al*, 1993; Efstratiadis, 1998). The *Zdbf2*-KO phenotype thus resembles an attenuated *Igf-1* deficiency, in agreement with half reduction but not total lack of circulating IGF-1. Interestingly, similar IGF-1-related growth defects have been reported upon alteration of other imprinted loci: the *Dlk1-Dio3* cluster and the *Cdkn1c* gene (Andrews *et al*, 2007; Charalambous *et al*, 2014). Finally, decreased *ZDBF2* levels have recently been associated with intra-uterine growth restriction (IUGR) in humans (Monteagudo-Sánchez *et al*, 2019). Whether this is linked to impaired fetal IGF-1 production would be interesting to assess.

IGF-1 secretion has been shown to drop in response to starvation, leading to disturbed growth physiology (Savage, 2013). Our findings support that limited food intake is probably the primary defect in *Zdbf2* deficiency, leading to IGF-I insufficiency in the critical period of postnatal development and consequently, growth restriction. First, we showed that *Zdbf2* is expressed in hypothalamic regions that contain neurons with functions in appetite and food intake regulation, such as the arcuate and paraventricular nucleus. Second, *Zdbf2-KO* neonates show hypothalamic downregulation of the *Npy* and *Agrp* genes that encode for orexigenic neuropeptides (Stanley & Leibowitz, 1984; Ollmann *et al*, 1997). These are likely direct effects, as the rest of the hypothalamic transcriptome is scarcely modified in *Zdbf2*-KO pups. Quantified changes in RNA-seq were not of large magnitude (although significant for *Npy*), but AgRP/NPY neurons represent a very small population of cells, present in the arcuate nucleus of the hypothalamus only (Andermann & Lowell, 2017). We are likely at the limit of detection when analyzing these genes in the whole hypothalamus transcriptome. Finally, *Npy* and *Agrp* downregulation was observed as early as 3 days after birth, along with a phenotype of reduced milk consumption, as measured by stomach weighing. Given our observations, we propose that ZDBF2 activates specialized hypothalamic neurons that motivate neonates to actively demand food (milk) from the mother right at birth, promoting the transition to oral feeding after a period of passive food supply *in utero*. This function agrees with the co-adaptation theory according to which genomic imprinting evolved to coordinate interactions between the offspring and the mother (Wolf & Hager, 2006). Despite considerable effort, we were unable to specifically detect the ZDBF2 protein with antibodies or using epitope-tagging approaches of the endogenous gene; future studies hopefully will bring clarity to which molecular function ZDBF2 carries in the hypothalamus.

In conclusion, we reveal here that decreasing *Zdbf2* compromises resource acquisition and body weight gain right after birth. Restricted postnatal growth can have a strong causal effect on metabolic phenotypes, increasing the risk of developing obesity in later life. This is observed in mouse KO models of the imprinted *Magel2* gene that maps to the Prader Willi syndrome region, and in PWS children themselves, who after a failure to thrive as young infants exhibit a catch-up phase leading to overweight and hyperphagia (Bischof *et al*, 2007). *Zdbf2*-KO mice do present a post-puberal spurt of growth, which attenuates their smaller body phenotype, but we never observed excessive weight gain, even after 18 months. It would be interesting to test whether the restricted postnatal growth of *Zdbf2*-KO mice may nonetheless increase the likelihood of metabolic complications when challenged with high-fat or high sugar diet.

## MATERIALS AND METHODS

### Mice

Mice were hosted on a 12 h/12 h light/dark cycle with free access to food and water in the pathogen-free Animal Care Facility of the Institut Curie (agreement number: C 75-05-18). All experimentation was approved by the Institut Curie Animal Care and Use Committee and adhered to European and National Regulation for the Protection of Vertebrate Animals used for Experimental and other Scientific Purposes (Directive 86/609 and 2010/63). For tissue and embryo collection, euthanasia was performed by cervical dislocation. The *Zdbf2*-KO and *Zdbf2*-GOF mutant mice lines were derived by CRISPR/Cas9 engineering in one-cell stage embryos as previously described (Greenberg *et al*, 2017), using two deletion-promoting sgRNAs. Zygote injection of the CRISPR/Cas9 system was performed by the Transgenesis Platform of the Institut Curie. Eight week-old superovulated C57BL/6J females were mated to stud males of the same background. Cytoplasmic injection of Cas9 mRNA and sgRNAs (100 and 50 ng/ul, respectively) was performed in zygotes collected in M2 medium (Sigma) at E0.5, with well-recognized pronuclei. Injected embryos were cultured in M16 medium (Sigma) at 37°C under 5% CO2, until transfer at the 1-cell stage the same day or at the 2-cell stage the following day in the infudibulum of the oviduct of pseudogestant CD1 females at E0.5. The founder mice were then genotyped and two independent founders with the expected deletion were backcrossed to segregate out undesired genetic events, with a systematic breeding scheme of *Zdbf2*-KO heterozygous females x WT C57Bl6/J males and *Zdbf2*-GOF heterozygous males x WT C57Bl6/J females to promote silent passing of the deletion. Cohorts of female and male N3 animals were then mated with WT C57Bl6/J to study the maternal and paternal transmission of the mutation.

The LacZ-*Zdbf2* reporter mouse line was derived from mouse embryonic stem (ES) cells from the European Conditional Mouse Mutagenesis Program (EUCOMM Project Number: *Zdbf2*_82543). Proper insertion of the LacZ construct was confirmed by long range PCR. However, upon Sanger sequencing, we found that the loxP site in the middle position was mutated (A to G transition at position 16 of the loxP site) in the original ES cells (Figure S4A). Chimeric mice were generated through blastocyst injection by the Institut Curie Transgenesis platform. We studied animals with an intact LacZ-KI allele, without FRT- or CRE-induced deletions.

### DNA methylation analyses

Genomic DNA from adult tissues was obtained following overnight lysis at 50°C (100mM Tris pH 8, 5mM EDTA, 200mM NaCl, 0.2% SDS and Proteinase K). DNA was recovered by a standard phenol/choloroform/isoamyl alcohol extraction and resuspended in water. Bisulfite conversion was performed on 0.5-1ug of DNA using the EpiTect Bisulfite Kit (Qiagen). Bisulfite-treated DNA was PCR amplified and either cloned and sequenced, or analyzed by pyrosequencing. For the former, 20-30 clones were Sanger sequenced and analyzed with BiQ Analyzer software (Bock *et al*, 2005). Pyrosequencing was performed on the PyroMark Q24 (Qiagen) according to the manufacturer’s instructions, and results were analyzed with the associated software.

### RNA expression analyses

Total RNA was extracted using Trizol (Life Technologies). To generate cDNA, 1ug of Trizol-extracted total RNA was DNase-treated (Ambion), then reverse transcribed with SuperscriptIII (Life Technologies) primed with random hexamers. RT-qPCR was performed using the SYBR Green Master Mix on the ViiA7 Real-Time PCR System (Thermo Fisher Scientific). Relative expression levels were normalized to the geometric mean of the Ct for housekeeping genes *Rrm2*, *B-actin* and/or *Rplp0*, with the ∆∆Ct method. Primers used are listed in Supplemental Table 2.

For RNA-sequencing, hypothalamus of three animals at 3dpp and two animals at 10dpp were collected for each genotype and RNA was extracted. Trizol-extracted total RNA was DNase-treated with the Qiagen RNase-Free DNase set, quantified using Qubit Fluorometric Quantitation (Thermo Fisher Scientific) and checked for integrity using Tapstation. RNA-seq libraries were cloned using TruSeq Stranded mRNA LT Sample Kit on total RNA (Illumina) and sequencing was performed in 100bp paired-end reads run on a NovaSeq (Illumina) at the NGS platform of the Institut Curie.

### LacZ staining

Whole brains and pituitary glands were fixed in 4% paraformaldehyde (PFA) in PBS (pH 7.2) overnight and washed in PBS. For sections, tissues were then incubated in sucrose gradients and embedded in OCT for conservation at −80°C before cryosectioning. Sections were first fixed 10 minutes in solution of Glutaraldehyde (0.02% Glutaraldehyde, 2mM MgCl2 in PBS). Tissues and sections were washed in washing solution (2mM MgCl2, 0.02% NP40, 0.01% C24H39NaO4 in PBS) and finally incubated at room temperature overnight in X- gal solution (5mM K4Fe(CN)63H20, 5mM K3Fe(CN)6, 25mg/mL X-gal in wash solution). After several PBS washes, sections were mounted with an aqueous media and conserved at room temperature before imaging.

### Histological analysis and RNA *in situ* hybridization

Embryos at E13.5 and E15.5 were embedded in paraffin and sectioned at a thickness of 5 μm. For histological analysis, paraffin sections were stained with Haematoxylin and Eosin. RNA *in situ* hybridization analysis on paraffin sections was performed following a standard procedure with digoxigenin-labeled antisense riboprobes as previously described (Gaston-Massuet *et al*, 2008).The antisense riboprobes used in this study [α-Gsu, *Pomc1*, *Lhx3*, *Pitx1, Avp, oxytocin and Ghrh*] have been previously described (Gaston-Massuet *et al*, 2008, 2016).

### Immunohistochemistry on histological sections

Embryos were fixed in 4% PFA and processed for immuno- detection as previously described (Andoniadou *et al*, 2013). Detection of hormones was carried out using antibodies for α-GH (rabbit polyclonal, NHPP AFP-5641801, 1:1000), α-TSH (rabbit polyclonal, NHPP AFP-1274789, 1:1000), α-PRL (rabbit polyclonal, NHPP AFP-425-10-91, 1:1000), α-ACTH (mouse monoclonal, 10C-CR1096M1, 1:1000) and α-LH (rabbit polyclonal, NHPP AFP-C697071P, 1:500).

### LUMINEX ELISA assay

Plasma from blood was collected in the morning at fixed time on EDTA from 5 and 15 day-old mice after euthanasia and stored at −20C until use. Samples were run in duplicates using the Mouse Magnetic Luminex Assay for IGF-1 (R&D System) and Milliplex Mouse Pituitary Magnetic Assay for GH (Merck) according to manufacturers’ instructions. Values were read on Bio-Plex® 200 (Bio-Rad) and analyzed with the Bio-Plex Manager Software.

### Phenotypic analyses of weight

Postnatal weight measurements were performed every two days from 1dpp to weaning age (21 days). Then, mice were separated according to their genotype, hosted in equal number per cage (*n*=5 to 6) and weighted once per week. As body weight is a continuous variable, we used a formula derived from the formula for the *t*-test to compute the minimum sample size per genotype: n=1+C(s/d)2, where *C* is dependent on values chosen for significance level (α) and power (1-β), *s* is the standard deviation and *d* the expected difference in means. Using α =5%, β=90% and an expected difference in means of 1g at 2 weeks and 2g later on, we predicted a minimum *n* of between 10 and 20 and thus decided to increase this number using *n* size between 20 and 30 per genotype, depending of the age and the sex of the mice. No animals were excluded from the analysis. Animals were blindly weighed until genotyping at weaning age. E18.5 embryos and post-natal organs were collected and individually, rinsed in PBS and weighted on a 0.001g scale. All data were generated using three independent mating pairs.

### Phenotypic analyses of post-natal lethality

Material for genotyping was taken the first day of birth and then number of pups for a given litter was assessed every day. To avoid bias, we excluded from the analysis the litters where all the pups died due to neglecting mothers, not taking care of their pups.

### DEXA scan analyses

The DEXA analysis allows the assessment of fat and lean mass, bone area, bone mineral content and bone mineral density. Practically, mutant and WT littermate males at 2 weeks were sent from the Animal Facility of Institut Curie to the Mouse Clinics along with their mother (Ilkirch, France). The phenotypic DEXA analysis was performed at 7 weeks of age using an *Ultrafocus DXA* digital radiography system, after the mandatory 5 week-quarantine period in the new animal facility. Mice were anesthetized prior analysis and scarified directly after measurements, without a waking up phase.

### RNA-seq data analysis

Adapters sequences were trimmed using TrimGalore v0.6.2 (https://github.com/FelixKrueger/TrimGalore). Paired-end reads were mapped using STAR_2.6.1a (Dobin *et al*, 2012) allowing 4% of mismatches. Gencode vM13 annotation was used to quantify gene expression using quantification mode from STAR. Normalization and differential gene expression were performed using EdgeR R package (v3.22.3)(Robinson *et al*, 2009). Genes were called as differentially expression if the fold discovery rate (FDR) is lower than 10%.

### Statistical Analyses

Significance of obtained data was determined by performing two-tailed unpaired, nonparametric Mann Whitney *t*- tests using GraphPad Prism6 software. *p* values were considered as significant when *p*≤0.05. Data points are denoted by stars based on their significance: ****: *p*≤0.0001; ***: p≤0.001; **: *p*≤0.01; *: *p*≤0.05.

## Data availability

The datasets produced in this study are available in the following databases: RNA-Seq data: Gene Expression Omnibus GSE153265 (https://www.ncbi.nlm.nih.gov/geo/query/acc.cgi?acc=GSE153265).

## ACKNOWLEDGEMENTS

We would like to thank members of the Bourc’his laboratory for continuous support and stimulation, M. Greenberg for critical reading of the manuscript, M. Charalambous for precious methodological and conceptual advices, T. Chelmicki and L. Marion-Poll for technical help and F. El Marjou (Transgenesis Platform of the Institut Curie Animal Facility) for CRISPR *in vivo* engineering. Luminex assays were performed by the LUMINEX Platform from the CHU Clermont-Ferrand. The laboratory of D.B. is part of the Laboratory d’Excellence LABEX (LABEX) entitled DEEP (11-LBX0044). This research was supported by the ERC (grant ERC-Cog EpiRepro) and the Bettencourt Schueller Foundation. C.G.M. and A.G. were supported by grants from Action Medical Research (GN2272) and BTL Charity (GN417/2238). J.G. was a recipient of PhD fellowships from DIM Biothérapies, Ile-de-France and La Ligue contre le Cancer.

## AUTHOR CONTRIBUTION

Conceptualization: D.B. and J.G; Methodology: D.B., J.G. and C.G.M. Formal analysis: J.G. and A.T.; Investigations: J.G., J.I., M.B., M.M., C.J., A.G.; Writing: J.G. and D.B.; Supervision: D.B.; Funding acquisition: D.B. and C.G.M.

## SUPPLEMENTAL DATA

### Generation and characterization of the *Zdbf2*-GOF line

We previously reported the CRISPR/Cas9-mediated generation of *Liz*-LOF mutant mice through targeting a 1.7kb deletion of the entire first exon of *Liz,* including its canonical transcription start site (TSS) (Greenberg et al., 2017). Among the founder mice, some carried a deletion smaller than expected, resulting from non-homologous end joining (NHEJ) repair of the cut induced by the left sgRNA only. Two of these lines –CRISPR-*Liz*_379 and CRISPR-*Liz*_397– exhibited a 924bp and 768bp deletion, respectively, which spanned the region upstream of the *Liz* TSS and left most of the exon 1 of *Liz* intact (Figure S4D). These two lines exhibited an unexpected pattern of DNA methylation at the *Zdbf2* locus upon maternal transmission only: heterozygous embryos at E9.5 with a maternally inherited allele showed a significant gain of sDMR DNA methylation as compared to their WT littermates (Figure S4G). We further demonstrated that the sDMR remained constitutively hypermethylated in adult tissues representative of the three germ layers (Figure S4E).

We previously demonstrated that sDMR DNA methylation is positively associated with expression *in cis* of *Zdbf2* in the postnatal hypothalamo-pituitary axis (Greenberg et al., 2017). To decipher if hypermethylation of the sDMR would be able to impact on *Zdbf2* expression, we measured by RT-qPCR the level of *Zdbf2* mRNA in the hypothalamus and the pituitary gland of 3 week-old animals. Focusing on the CRISPR-*Liz*_379 line, we showed increased *Zdbf2* expression upon maternal transmission of the deletion (Figure S4F) and thus named this mutant as *Zdbf2*-gain-of-function (*Zdbf2*-GOF). Allelic analysis from hybrid mice between C57Bl6/J and JF1 strains demonstrated that this gain-of-function was due to ectopic DNA methylation at the sDMR on the maternal allele and ectopic expression of the normally silent maternal *Zdbf2* allele (Figure 3A and B). Reactivation of the maternal allele resulted in bi-allelic, rather than strictly paternal mono-allelic expression of *Zdbf2*. The level of expression of the maternal allele was not equal to the expression of the paternal allele, resulting in a net 1.7-fold increase of *Zdbf2* in *Liz*-GOF animals compared to WT levels (Figure S4F). This was linked to a mosaic gain of DNA methylation on the maternal allele of *Zdbf2*-GOF animals, with 70% versus 90-95% of paternal methylation levels. Allelic analysis revealed cellular heterogeneity in the acquisition of methylation of the maternal allele (Figure S4H). In contrast to the effect of the maternal allele, paternal transmission of the deletion behaved as a silent mutation (Figure S4E and F).

In sum, we have generated mutant mice with loss-of-imprinting of the *Zdbf2* locus, harboring increased *Zdbf2* dosage in the postnatal brain, as a result of abnormal biallelic expression. The deletion had no effect on the sex and Mendelian distributions upon both maternal and paternal transmission (Figure S5A and B) and the mice survived at the same rate as their WT littermates (Figure S5C). Moreover, we did not detect any fertility or behavioral phenotype in the *Zdbf2*-GOF mice.

**Figure S1:**
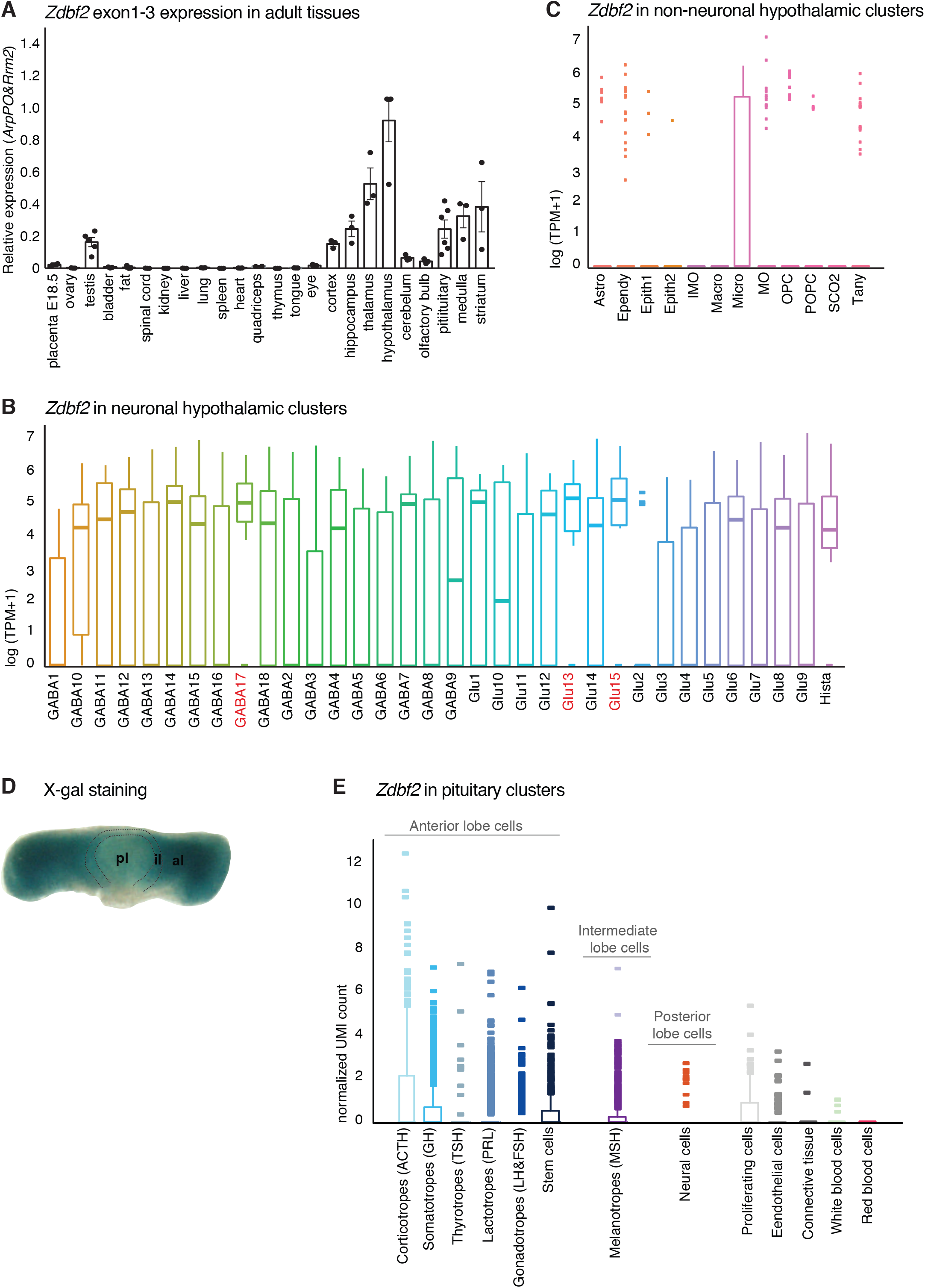
*Zdbf2* expression from pituitary and hypothalamus. (**A**) Expression of *Zdbf2* measured by RT-qPCR in a bank of adult (6-weeks old) mouse tissues. *Zdbf2* is preferentially expressed in the brain and hypothalamus is the tissue showing the highest level of expression. Data are shown as means ± s.e.m. This figure has been adapted from Greenberg et *al* 2017. (**B, C**) Expression of *Zdbf2* in hypothalamic neuronal (**B**) and non-neuronal (**C**) cell clusters. Clusters where *Zdbf2* expression is the highest are highlighted in red. Single-cell RNA-seq datasets from the hypothalamus of 8 to 10 week-old B6D2F1 females from Chen *et al.*, Cell Reports, 2017. (**D**) Whole-mount pituitary from a *Zdbf2*-LacZ KI mouse stained with X-gal. pl, posterior lobe; il, intermediate lobe; al, anterior lobe. (**E**) Expression of *Zdbf2* in pituitary cell clusters. *Zdbf2* expression is found predominantly in cluster of cells from the anterior and intermediate lobes. Single-cell RNA-seq datasets from pituitaries of 7 week-old C57BL/6J males from Cheung *et al.*, Endocrinology, 2018.

**Figure S2:**
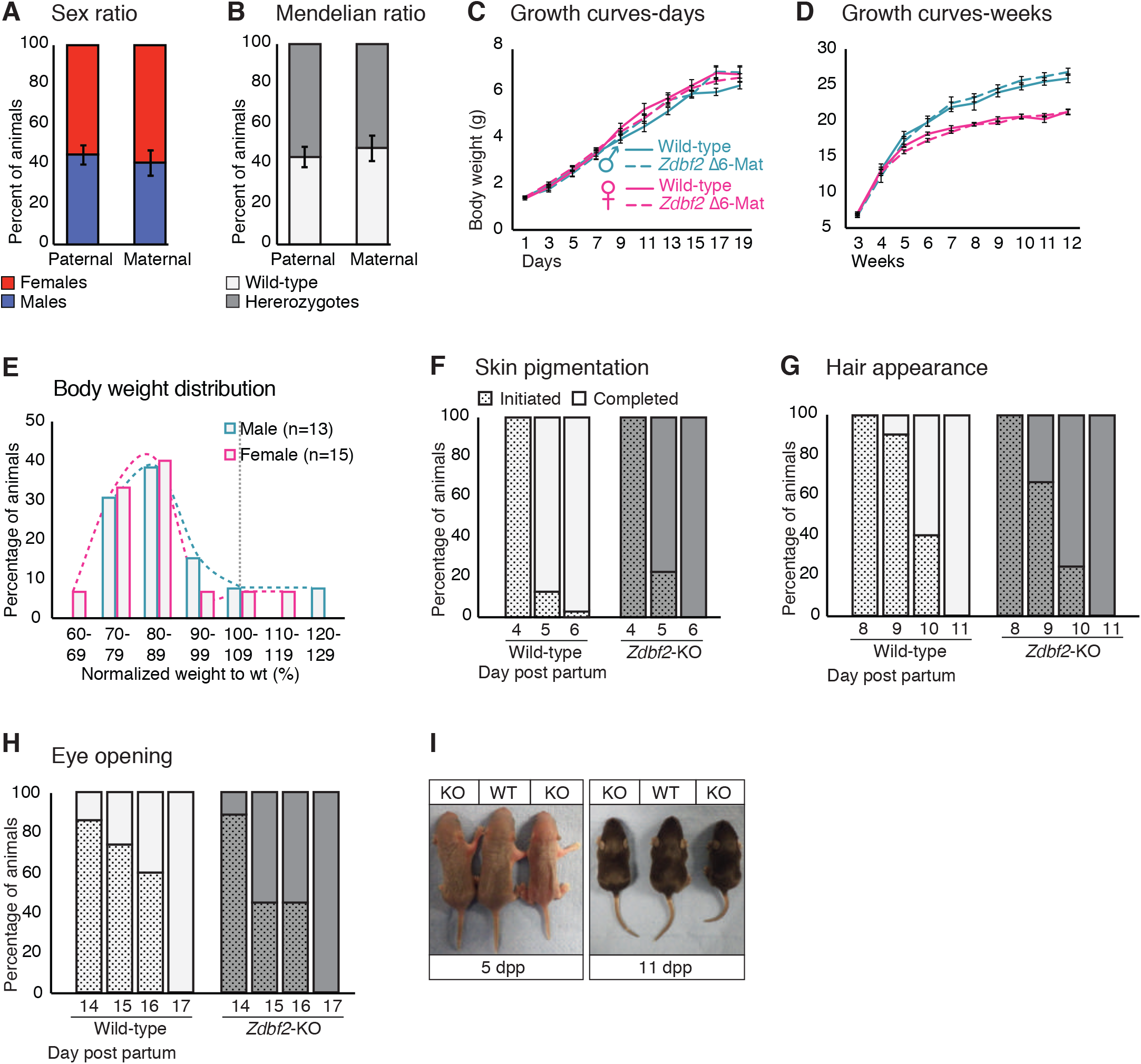
*Zdbf2*-KO pups acquire normal hallmarks of postnatal development. (**A, B**) Sex ratio (**A**) and genotype ratio (**B**) observed at birth (1dpp) among *Zdbf2* heterozygotes from paternal and maternal transmission of the deletion after three backcrosses. Data are shown as means ± s.e.m. from individuals from n= 32 and 12 litters from paternal and maternal transmission respectively. (**C, D**) Growth curves as in Figure 2C and Figure 2D, respectively, but comparing mutants with a maternal transmission of the Zdbf2 deletion with their WT littermates. (**E**) Frequency of body weight distribution for normalized weights of 2 week-old *Zdbf2*-KO mice relative to WT mice (100%) grouped into 10% bins. The weight reduction phenotype appears to be highly penetrant. (**F-H**) Graphical representation of the timing of three typical hallmarks of postnatal development: skin pigmentation (**F**), hair appearance and eye opening (**H**) between WT and *Zdbf2*-KO littermates. (**I**) Representative photography of WT and *Zdbf2*-KO female littermates at 5dpp (left) and 11dpp (right).

**Figure S3:**
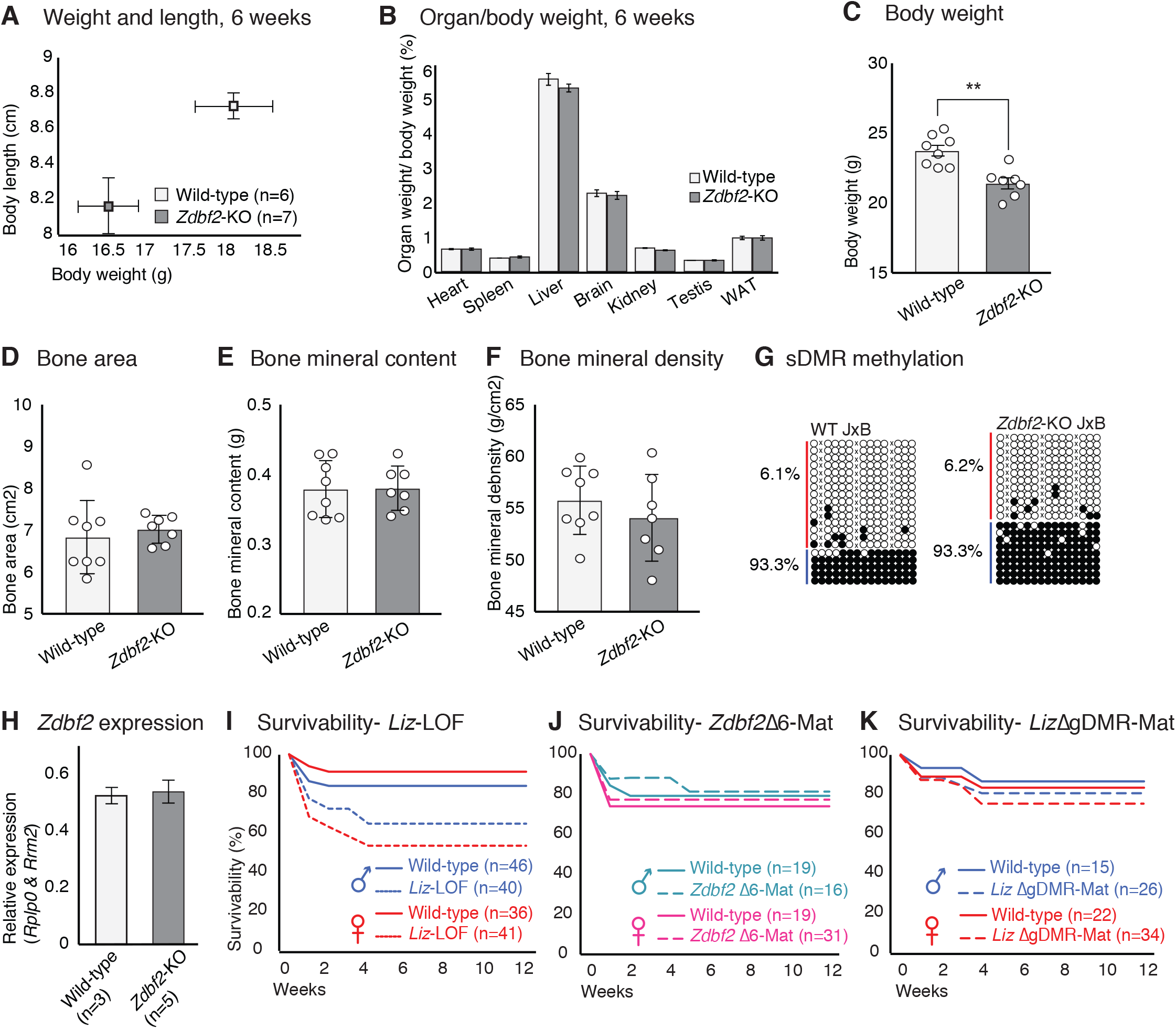
Phenotypic and molecular characterization of *Zdbf2* mutants. Average body length and weight of n WT and *Zdbf2*-KO mice at 6 weeks. (**B**) Individual organ weight plotted as percentage of total body weight in *Zdbf2*-KO mice and WT littermates at 6 weeks. (**C-F**) Dual-energy X-ray absorptiometry (DXA) showing calculation of body weight (**C**), bone area (**D**), bone mineral content (**E**) and bone mineral density (**F**). Data are shown as means [)5] s.e.m from n=8 WT and 7 *Zdbf2*-KO males. (**G**) Bisulfite cloning and sequencing of the sDMR from hypothalamus DNA of 3 week-old hybrid WT (top) and *Zdbf2*-KO (bottom) mice (JF1 x *Zdbf2*-KO cross) reveals that the deposition of DNA methylation at the sDMR is not affected by *Zdbf2* deletion. Red, maternal alleles; blue, paternal alleles. cross, informative JF1 SNP. (**H**) RT-qPCR measuring Z*dbf2* expression from exon 1 to 3 in 3 week-old hypothalamus reveals that *Zdbf2* transcriptional output is not affected by the deletion of the exon 6. Statistical analyses were performed by a two-tailed, unpaired, nonparametric Mann Whitney t test. **p<0.01. (**I-K**) Kaplan-Meier curves of the survivability as in Figure 2I but comparing WT and *Liz*-LOF males and females (**I**), WT and *Zdbf2*-∆exon6 males and females upon maternal transmission of the deletion (**J**) and WT and *Liz*-∆ gDMR males and females upon maternal transmission of the deletion (**K**).

**Figure S4:**
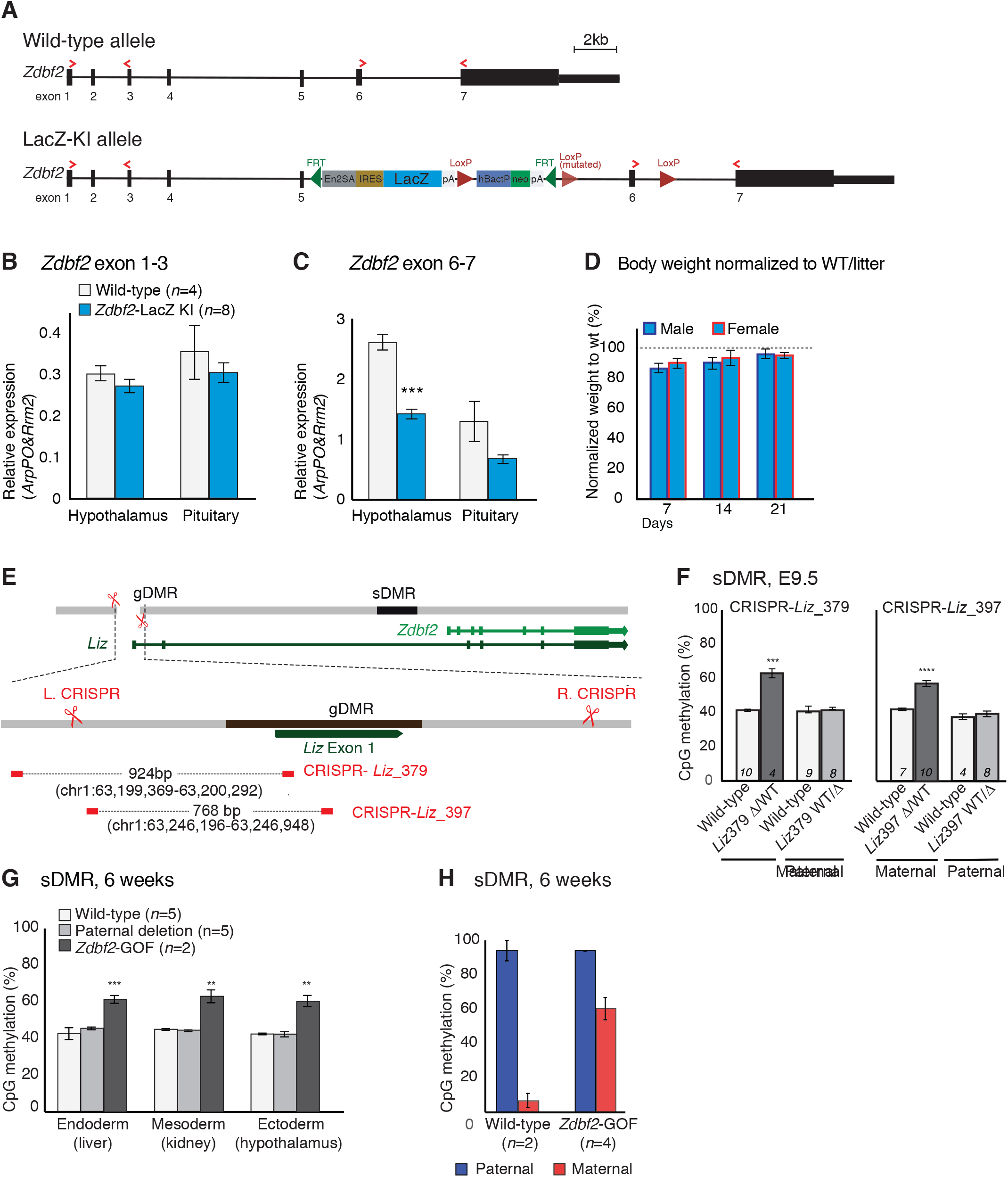
Characterization of partial *Zdbf2*-LOF and *Zdbf2*-GOF mouse lines. (**A**) Scheme of the wild-type and LacZ-KI *Zdbf2* alleles. The seven exons of the *Zdbf2* transcript are represented by black boxes and the primers used for the RT-qPCR in (**B,C**) are labeled as red arrows. The LacZ-KI allele contains an insertion of: a trapping cassette (En2SA-IRES-LacZ-pA), a Neo cassette under the human beta actin promoter (hbactP-Neo-pA) together with FRT sites to delete the cassettes and LoxP sites to delete exon 6. However, we found the middle loxP site to be non-functional because of a point mutation in the original EUCOMM-provided ES cells. (**B, C**) *Zdbf2* expression measured by RT-qPCR from exon 1-3 (**B**) and exon 6-7 (**C**) in hypothalamus and pituitary of WT and *Zdbf2*-LacZ KI individuals. The insertion schematized in (**A**) leads to a ~ 2-fold decrease in *Zdbf2* expression at exon 6-7, probably due to exon skipping. Data are shown as means ± s.e.m. from n individuals. Statistical analyses were performed by a two-tailed, unpaired, nonparametric Mann Whitney t test. ***p<0.001. (**D**) Normalized body growth of *Zdbf2*-LacZ KI mice to their WT littermates (100%) followed at 1, 2 and 3 weeks shows a slight decrease in weight of both males and females. Data are shown as means ± s.e.m. from 8 litters. (**E**) Graphic model of the ∆gDMR CRISPR deletion leading to the generation of the *Liz*_379 founder line (*Zdbf2*-GOF). Deletion is the results of a single cut by Cas9 at the left sgRNA position, followed by NHEJ. (**F**) Bisulfite-pyrosequencing from post-implantation embryos at E9.5 shows increase of sDMR CpG methylation only upon maternal transmission of the deletion, both in *Liz*_379 (left) and *Liz*_397 (right) CRISPR lines. (**G**) Bisulfite-pyrosequencing shows increased sDMR methylation in tissues from 6 week-old mice when the deletion is maternally transmitted. The paternal transmission of the deletion behaves as silent. (**H**) Quantification of allelic sDMR CpG methylation measured by bisulfite cloning and sequencing from n WT and *Zdbf2*-GOF adult tissues. The gain of sDMR methylation from the maternal allele in the mutant compared to their WT littermates is about 65%.

**Figure S5:**
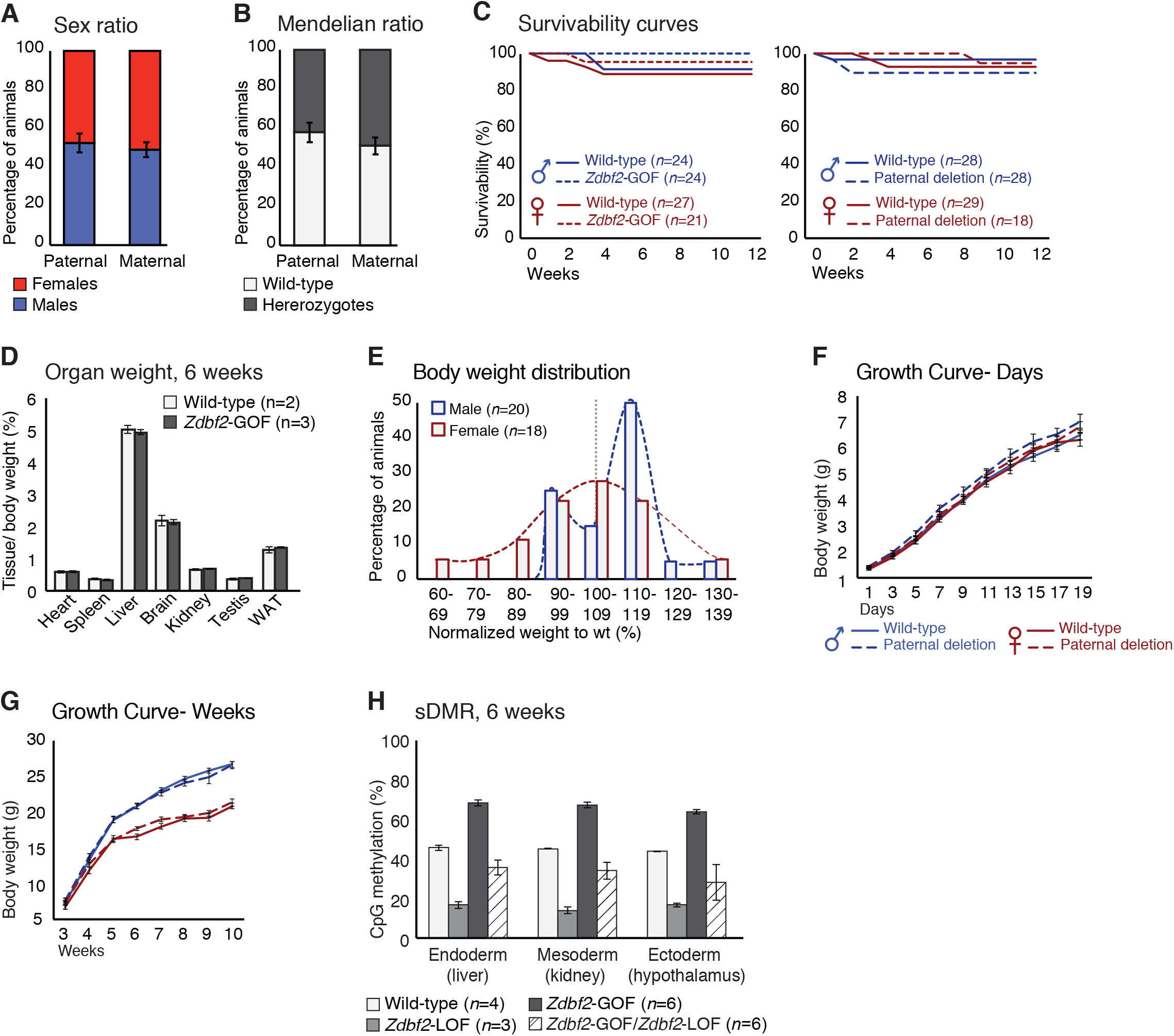
Phenotypic characterization of *Zdbf2*-GOF mutants. (**A, B**) Sex ratio (**A**) and genotype ratio (**B**) observed at birth among heterozygotes upon paternal and maternal transmission of the *Zdbf2*-GOF mutation after three backcrosses of the *Liz*_379 line. Data are shown as means ± s.e.m. from individuals from n= 15 and 20 litters from paternal and maternal transmission respectively. (**C**) Kaplan-Meier curves of the survivability from birth to 12 weeks showing no lethality phenotype upon maternal (left) and paternal (right) transmission of the deletion, both in males and females. Data are shown as means ± s.e.m. from n individuals. (**D**) Individual organ weight plotted as percentage of total body weight in *Zdbf2*-GOF mice and WT littermates at 6 weeks. (**E**) Frequency of weight distribution for normalized weights of 2-week-old *Zdbf2*-GOF mice relative to WT mice (100%) grouped into 10% bins. The male specific overgrowth phenotype appears to be penetrant. Females show a normal distribution of weight, confirming the sex-specific phenotype seen in C and D. (**F, G**) Growth curves as in Figure 5D and 5E, but comparing the mutants with a paternal transmission of the deletion with their WT littermates. The paternal deletion behaves as a silent mutation with no growth-related phenotype in the progenies. n=10–25 mice were analyzed per genotype, depending on age and sex. (**H**) Bisulfite-pyrosequencing shows an almost complete rescue of the *Zdbf2*-LOF defects of sDMR methylation in tissues of *Zdbf2*-GOF /*Zdbf2*-LOF mice at 6 weeks. Data are shown as means ± s.e.m. from n individuals. Statistical analyses were performed by a two-tailed, unpaired, nonparametric Mann Whitney t test. *p ≤ 0.05

**Figure S6:**
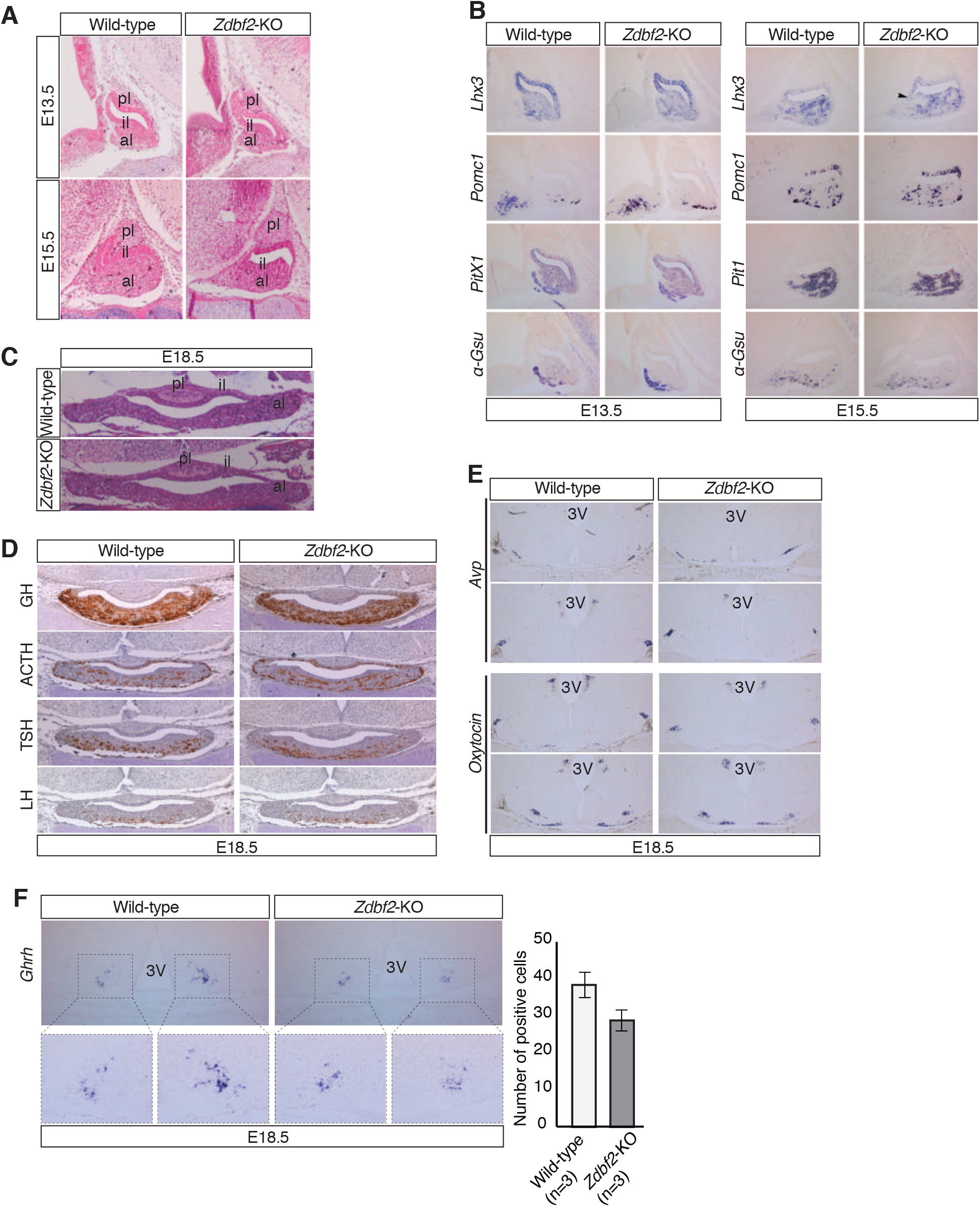
Normal development of the hypothalamo-pituitary axis in *Zdbf2*-KO mice. (**A**) Haematoxylin-Eosin (H&E) staining reveals normal development of the pituitary at embryonic day (E) 13.5 and E15.5 embryonic days. (**B**) RNA in situ hybridization for pituitary lineage-specific genes reveals normal mRNA expression of cell specification markers in the pituitary of the *Zdbf2*-KO embryos at E13.5 and E15.5. (**C**) Morphological assessment of the mature E18.5 pituitary gland by H&E staining shows no phenotype in the pituitary gland before birth. (**D**) Immunohistochemistry analysis of four pituitary hormones (GH, ACTH, TSL, LH) at E18.5 suggests no obvious defect in hormone-producing cells in the anterior lobe of the pituitary before birth. (**E**) In situ hybridization for two hypothalamic neuropeptides, arginine vasopressin (AVP) (top) and oxytocin (bottom), in the developing E18.5 hypothalamus suggests no apparent differences in AVP and oxytocin transcripts. (**F**) RNA in situ hybridization for *Ghrh* on E18.5 hypothalamus shows trend towards decreased expression of *Ghrh* in mutant embryos compared to controls, although difference is not statistical significant. Quantification (right panel) represents the means ± s.e.m of the number of *Ghrh* positive cells over n=3 independent experiments. Overall, results in **A-F** are congruent with the observation of normal weight distribution in *Zdbf2*-KO E18.5 embryos (Figure 2B). pl, posterior lobe; il, intermediate lobe; al, anterior lobe; 3V, third ventricle of the hypothalamus.

**Figure S7:**
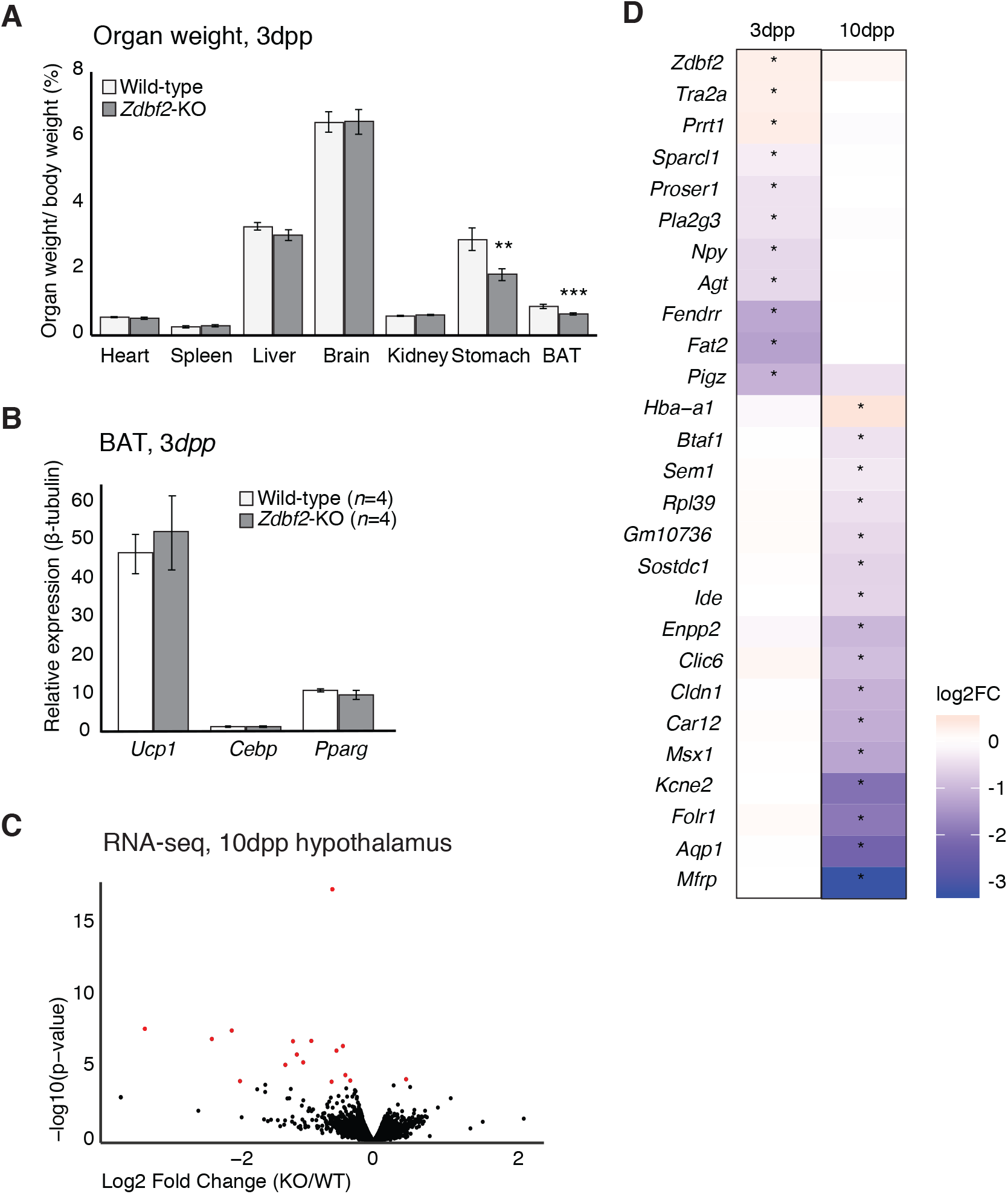
Characterization of the early postnatal feeding behavior in *Zdbf2*-KO pups. Individual organ weight plotted as percentage of total body weight in *Zdbf2*-KO mice and WT littermates at 3dpp. (**B**) RT-qPCR from BAT measuring the mRNAs level of three markers of BAT activity (*Ucp-1*, *Cebp* and *Pparg*) between WT and *Zdbf2*-KO at 3dpp. (**C**) Volcano plot representation of RNA-seq of 10dpp hypothalamus of *Zdbf2*-KO versus WT littermates. Red dots: differentially expressed genes with a threshold of FDR<10%. (**D**) Heatmap showing the log2 fold change of expression of the differentially expressed genes (DEGs) in the hypothalamus of *Zdbf2*-KO pups at 3 and 10dpp. There is no overlap between the DEGs at 3 and 10dpp. Stars represent significant change of expression (FDR<10%).

---

**Supplemental Table 1:** Survivability counts from different transmission of the *Zdbf2*-KO allele. (A) Number of living pups followed every two days from 1 to 21dpp and every week from 21 to 84dpp from 24 litters coming from [female WT x male *Zdbf2* KO/WT] crosses (paternal transmission). These 24 litters come from 4 different crosses including 8 females and 4 males. Day 1 corresponds to the day of birth (1dpp). We can observe that the number of living *Zdbf2*-KO (males and females) starts decreasing from 3dpp onwards. (B) Number of living pups followed every day from 1 to 21dpp from 19 litters coming from [female WT x male *Zdbf2* KO/KO] crosses (homozygote transmission). These 19 litters come from 5 different crosses including 10 females and 5 males. All pups from these crosses inherited a mutated allele from their dad and are thus *Zdbf2*-KO. (C) Number of living pups followed every week from 1 to 84dpp from 11 litters coming from [female *Zdbf2* KO/WT x male WT] crosses (maternal transmission, silent mutation). These 11 litters come from 2 different crosses including 4 females and 2 males. We did not observe any survivability bias within those litters.

| A. *Zdbf2*-KO (paternal transmission) |  |  |  |  |  |  |  |  |  |  |  |  |  |  |  |  |  |  |  |  |  |
| --- | --- | --- | --- | --- | --- | --- | --- | --- | --- | --- | --- | --- | --- | --- | --- | --- | --- | --- | --- | --- | --- |
| Days post partum |  | 1 | 3 | 5 | 7 | 9 | 11 | 13 | 15 | 17 | 19 | 21 | 28 | 35 | 42 | 49 | 56 | 63 | 70 | 77 | 84 |
| n=24 litters | Males WT | 44 | 37 | 37 | 37 | 37 | 37 | 37 | 37 | 37 | 36 | 36 | 36 | 36 | 36 | 36 | 36 | 36 | 36 | 36 | 36 |
|  | Female WT | 50 | 44 | 44 | 44 | 44 | 44 | 44 | 44 | 43 | 43 | 40 | 39 | 39 | 39 | 39 | 39 | 39 | 39 | 39 | 39 |
|  | Males KO | 30 | 17 | 17 | 17 | 17 | 16 | 16 | 16 | 16 | 16 | 16 | 16 | 16 | 16 | 16 | 16 | 16 | 16 | 16 | 16 |
|  | Female KO | 57 | 30 | 30 | 30 | 30 | 29 | 29 | 29 | 28 | 28 | 28 | 28 | 28 | 28 | 28 | 28 | 27 | 27 | 27 | 27 |

**Supplemental Table 2:**
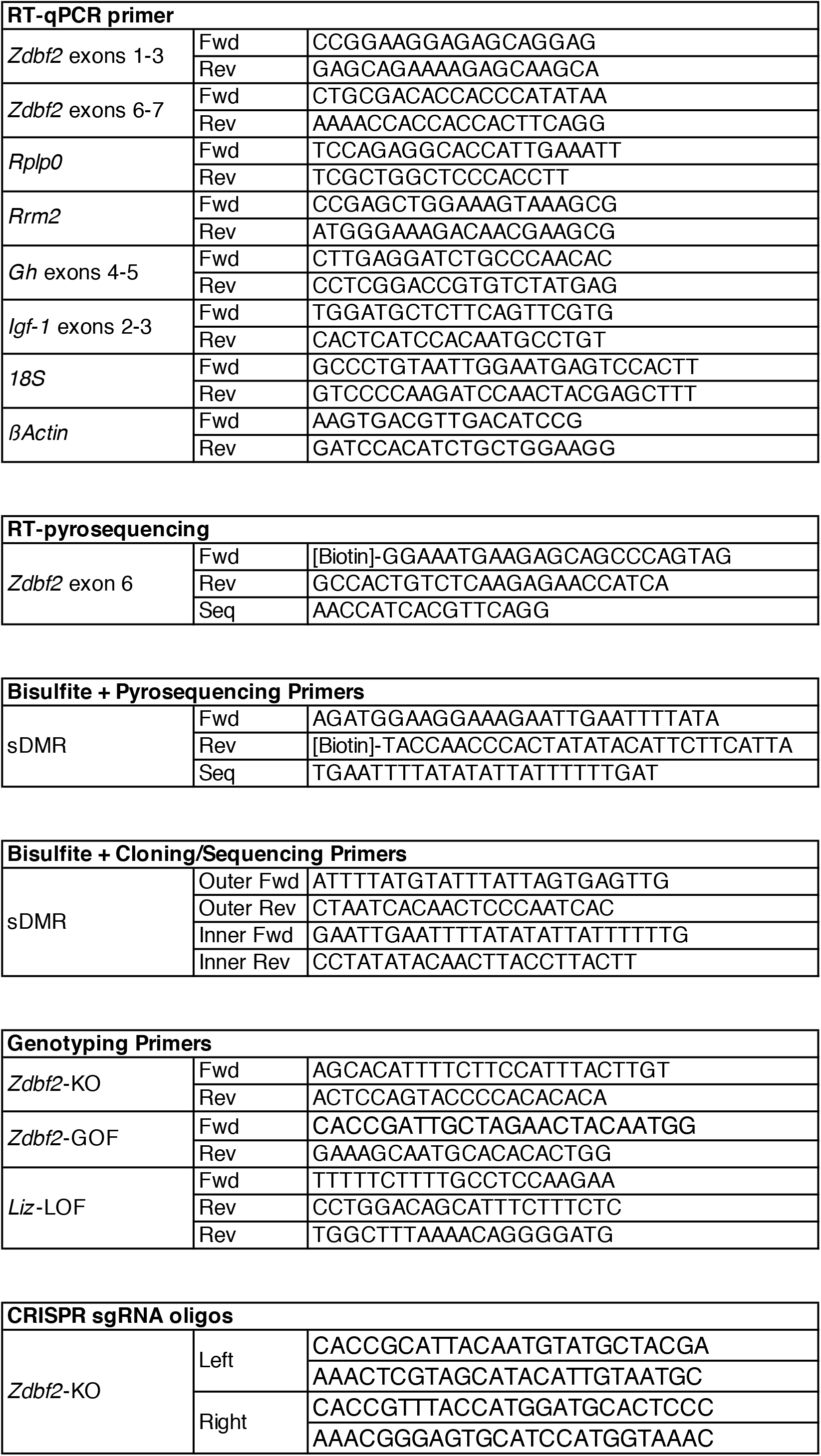
List of primers used in this study.

